# Accurate and robust 3D genome feature discovery from multiplexed DNA FISH

**DOI:** 10.1101/2025.11.26.690837

**Authors:** Hongyu Yu, Lingbo Zhou, Liangqi Xie, Caterina Strambio-De-Castillia, Ming Hu, Yun Li

## Abstract

Multiplexed DNA FISH (also known as FISH Omics) is a powerful imaging technology to map three-dimensional (3D) genome organization in thousands of single cells at kilobase resolution. However, existing computational approaches to identify 3D genome features, such as A/B compartments, topologically associating domains (TADs) and chromatin loops, have been mainly adapted from Hi-C and do not account for the distinct measurement errors across the three XYZ physical axes. Here we present ArcFISH (**a**ccurate and **r**obust **c**hromatin analysis from FISH data), which explicitly models axis-specific measurement errors in a unified statistical framework to detect multi-level 3D genome features. Comprehensive evaluations using multiple simulated as well as experimental datasets show that ArcFISH reliably identifies chromatin loops under heterogeneous measurement errors, detects TAD boundaries highly enriched for active histone marks H3K4me3 and H3K36me3, and requires as few as 30 chromatin traces to faithfully assign A/B compartments. Applying ArcFISH to multiplexed DNA FISH data from mouse embryonic stem cells and mouse excitatory neurons reveals cell-type-specific chromatin loops and TADs, overlapping with differentially expressed genes. ArcFISH is freely available at https://hyuyu104.github.io/ArcFISH.

## Introduction

The eukaryotic genome is not randomly organized in the nucleus^1,2^. Instead, chromatin forms complex three-dimensional (3D) structures^3–6^, which, ordered by scale, include chromosome territories, A/B compartments^7^, topologically associated domains (TADs)^8^, and long-range chromatin loops^9,10^. These hierarchical features are highly cell-type-specific and reveal functionally coordinated regions of the genome, thus playing a key role in gene regulation^11–13^. Genome-wide chromatin conformation capture (i.e., Hi-C) has been widely used to study 3D genome, with the typical procedure involving: chromatin fixation, fragmentation, cross-linking and then sequencing ^7,10,14^. This results in a contact matrix, where each entry records how frequently pairs of DNA segments interact with each other among the cell population. A variety of computational methods have been developed to identify A/B compartments, TADs, and loops from such matrices^15–20^.

Recent advances in imaging technologies such as DNA-MERFISH^21,22^, DNA-seqFISH+^23–25^, ORCA^26–28^, Hi-M^29,30^, or MINA^31^, provide an appealing alternative to Hi-C and Hi-C-derived sequencing-based approaches^32,33^. Instead of relying on the construction of two-dimensional contact matrices to infer 3D genome structures, imaging technologies enable *direct* visualization of the individual genome segments by sequential Fluorescence In-Situ Hybridization (FISH) segments. The resulting data, which we refer to as chromatin tracing data, contain the 3D coordinates of imaged genomic segments in each single cell.

However, despite the distinct data generation mechanisms, most existing computational pipelines for 3D genome feature identification from chromatin tracing data first construct a pairwise contact matrix and apply methods developed for Hi-C^34–37^. Specifically, the common practice is to calculate the pairwise Euclidean distance between different chromatin segments within each single cell, compute the average pairwise distance matrix across a population of cells, and adapt Hi-C-oriented computational methods to the average pairwise distance matrix^34–36,38^. This approach ignores the varying levels of measurement uncertainties that exists in chromatin tracing data at the level of each of the individual x, y, and z physical axes, leading to inaccurate 3D genome feature characterization.

Since images are acquired in x-y planes while moving along the z-axis, the x and y axes tend to share similar levels of measurement uncertainties, whereas the z-axis often exhibits a different level of uncertainty^22,23,27^. When the z-axis has a higher level of noise, the pairwise distance matrices are often dominated by this z-axis-specific noise, masking the more accurate information from the x and y axes. On the other hand, when the z-axis is measured more accurately than the x and y axes, current pairwise-distance-based computational methods cannot fully leverage the more accurate signal from the z-axis. After the 3D coordinates in chromatin tracing data are converted to pairwise distance matrices, no axis-level information is available as Euclidean distance is the sum of pairwise differences, unweighted across each axis. Therefore, developing an approach to process axes separately and to properly combine information from different axes has the potential to fully mine the rich information in the chromatin tracing data.

Here we introduce a model-based method that accounts for the distinct measurement error in each axis separately and integrates the results from different axes via adaptive weighting. We then describe how to apply this method to identify A/B compartments, TADs and chromatin loops from chromatin tracing data. We name our method ArcFISH (**a**ccurate and **r**obust **c**hromatin analysis from FISH data). By benchmarking against existing methods across a variety of chromatin tracing datasets, from both mice and humans, and across different imaging protocols, we show that ArcFISH consistently outperforms current pipelines in terms of accuracy, interpretability, and biological relevance. Application of ArcFISH to a mouse brain DNA seqFISH+ dataset revealed highly cell-type-specific TADs and chromatin loops. We found that differential TAD boundaries between mouse excitatory neurons and mouse embryonic stem cells potentially correspond to differentially expressed genes, and we also observed increased loop activity in neurons compared with embryonic stem cells in the imaging region. Our ArcFISH software is written in Python, with full documentation available at https://hyuyu104.github.io/ArcFISH.

### Overview of ArcFISH

ArcFISH takes chromatin tracing data as input and outputs chromatin loops, TADs, and A/B compartments identified from the target imaging region. All three tasks are based on a four-step procedure: axis-specific weight estimation, filtering and normalization, task-dependent axis-wise processing, and weighted combination of processed results (Figure 1). First, to estimate the weight, we model each chromatin trace as a 3D Gaussian process, and the observed chromatin tracing data is this 3D Gaussian process plus an axis-specific random noise (see Methods). By assuming the underlying pairwise distance matrix is of low-rank (see Methods), we estimate the variance of the noises and use their inverse as axis weights, analogous to the inverse-variance estimator widely adopted in meta-analysis^39^. Next, we calculate the pairwise difference matrix for each axis, filter out potential mismeasurements, and normalize by 1D genomic distance as commonly performed in Hi-C analyses^19,40,41^.

**Figure 1.**
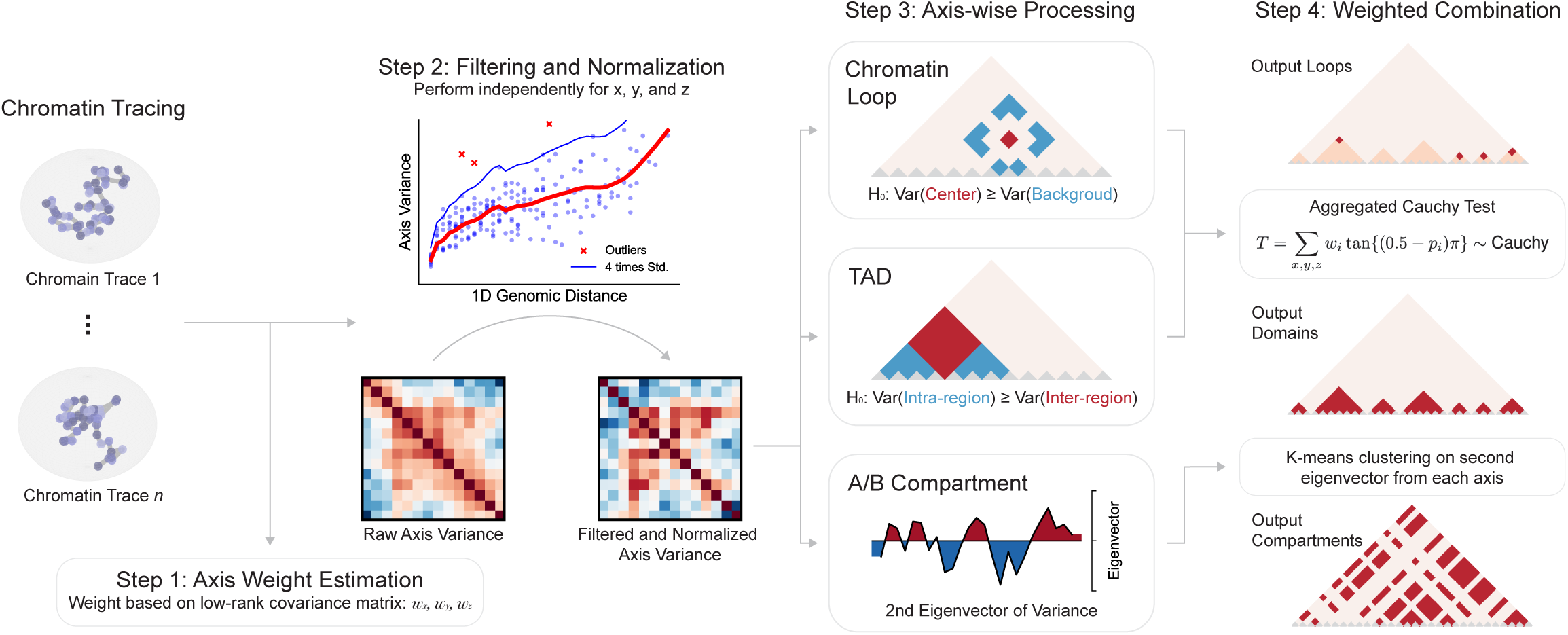
ArcFISH processes each axis separately to ensure robustness and accuracy. Input chromatin tracing data contains the 3D coordinates of each imaging segment in each chromatin trace (*n* traces in total) denoted by a node in Figure. ArcFISH uses raw data to estimate a weight for each axis based on the low rank assumption. In the second step, pairwise differences for each axis (*nxpx p* values if *p* is the number of imaged segments) are filtered based on a lowess curve against the 1D genomic distance and then normalized to have unit variance. Instead of storing all the pairwise differences, only a *px p* axis variance matrix (pairwise differences averaged across *n* traces) is stored. The third step uses the normalized variance matrices to perform task specific processing. For chromatin loops, ArcFISH tests whether the query entry has significantly smaller variance than the local background. For TADs, given a segment, ArcFISH compares the inter-domain variance and the intra-domain variance. If the intra-domain variance is significantly smaller, the segment is labeled as a TAD boundary. For A/B compartments, ArcFISH performs eigenvalue decomposition on each axis and keeps the second eigenvectors. In the last step, the test results for loops and TADs are combined by aggregated Cauchy test, accounting for axis weights. A/B compartments are determined by applying K-means clustering to the weighted second eigenvectors.

For axis-wise processing, we develop separate strategies for each task based on the normalized pairwise difference matrices (Figure 1). For loop calling, we test the variance of each entry in the pairwise difference matrix against the variances of its local background, defined as entries between 25 Kb to 50 Kb from the testing entry. The p-values for x, y, and z axes are then computed from a F-distribution based on the pairwise difference matrix from the respective axis. Mathematically, as squared Euclidean distance is the sum of variances of pairwise difference, small p-values effectively select entries with smaller pairwise distance compared to their local background, which corresponds to chromatin loops. For TAD calling, we consider a 200 Kb window surrounding each segment and test inter-domain variance against intra-domain variance. The p-values are computed using a F-test as well and large difference in variance signifies potential TAD boundaries analogous to insulation scores^42^. For A/B compartment assignments, eigenvalue decomposition is applied to the pairwise difference matrices, and the second eigenvectors from each axis are kept.

For the last step, we apply the weights to combine the axis-wise information obtained in the third step (Figure 1). For both loop and TAD results, we combine the p-values by the aggregated Cauchy test^43^ to obtain a single p-value for each segment pair or each segment and convert the p-values to false discovery rates (FDRs) to adjust for multiple testing. Segment pairs or segments with FDRs smaller than 0.1 are reported as chromatin loops or TAD boundaries, respectively. For A/B compartment calling, we multiply the eigenvectors by their corresponding weights and perform K-means clustering in this weighted feature space. This results in two clusters, and the cluster with higher transcription start sites (TSSs) density is assigned as the A compartment.

The ArcFISH software supports importing chromatin data in compliance with the 4DN FISH Omics Format for Chromatin Tracing (FOF-CT)^44^ data, which was recently developed by the 4DN^4^ consortium for the FAIR sharing of multiplexed FISH results^35,36,45^. The input data are stored as AnnData objects^46^ by chromosome (Figure S1). When performing filtering and normalization, since the storage complexity of pairwise difference matrices grows quadratically with the number of chromatin segments and linearly with the number of traces, we perform the computation by Dask arrays^47^ and divide pairwise difference matrices to 1GB chunks, thus allowing processing larger-than-memory matrices and parallel computation on servers. We also make some algebraic optimizations on test statistics so that all step 3 operations can be done on precomputed axis variance matrices, further minimizing computation cost (see Supplementary Methods Section 4). Finally, ArcFISH offers various visualization tools to facilitate easy interpretation.

### ArcFISH accurately and robustly identifies chromatin loops

We benchmarked our chromatin loop calling algorithm on mouse embryonic stem cells (mESCs) DNA seqFISH+ dataset^23^. The original data contain 60 consecutive segments on each chromosome at 25Kb resolution, resulting in an approximately ∼1.5Mb region in each of the 20 chromosomes. Except the X chromosome, which only has 483 chromatin traces, about 950 traces are generated for the other 19 chromosomes, with chromosome 19 having the most (1,045) and chromosome 15 having the least (897). To examine the effect of heterogeneous measurement noises, we first performed the experimental data-based simulation and generated 800 traces with uniform measurement errors in x, y, and z axes from the original data and added 0, 50, 100, and 200 nm noise to the z-axis (see Methods). We then applied both SnapFISH and ArcFISH to 10 replicates from each noise level and evaluated their recall and precision (Figure 2a and 2b). As a working ground truth, we collected six reference loop lists identified from mESCs: MAPS-identified loops from CTCF PLAC-seq data^48^, MAPS-identified loops from H3K4me3 PLAC-seq data^48^, CTCF ChIA-PET loops^49^, POLR2A ChIA-PET loops^49^, HiCExplorer^50^ and FitHiC2^51^ loops from bulk Hi-C^14^. The true positives in recall and precision are defined as loops present in at least two of the above six loop lists, resulting in a total of 154 loops (see Methods). As noise level increases, the average precision of SnapFISH drops from 34% to 23% (69% of zero noise) while the precision of ArcFISH changes slightly from 52% to 49% (95% of zero noise) (Figure 2c and S4a). Notably, the precision of ArcFISH is maintained with the consistent recall (Figure 2c). For a comparison, the average recall of SnapFISH at 150 nm noise is only 53% of its recall at 0 nm noise while this number is 98% for ArcFISH (Figure S4b). Therefore, as measurement uncertainties become more heterogeneous, traditional methods relying on pairwise distance and ignoring axis-specific measurement errors have worse performance, while ArcFISH captures more reliable loop signals.

**Figure 2.**
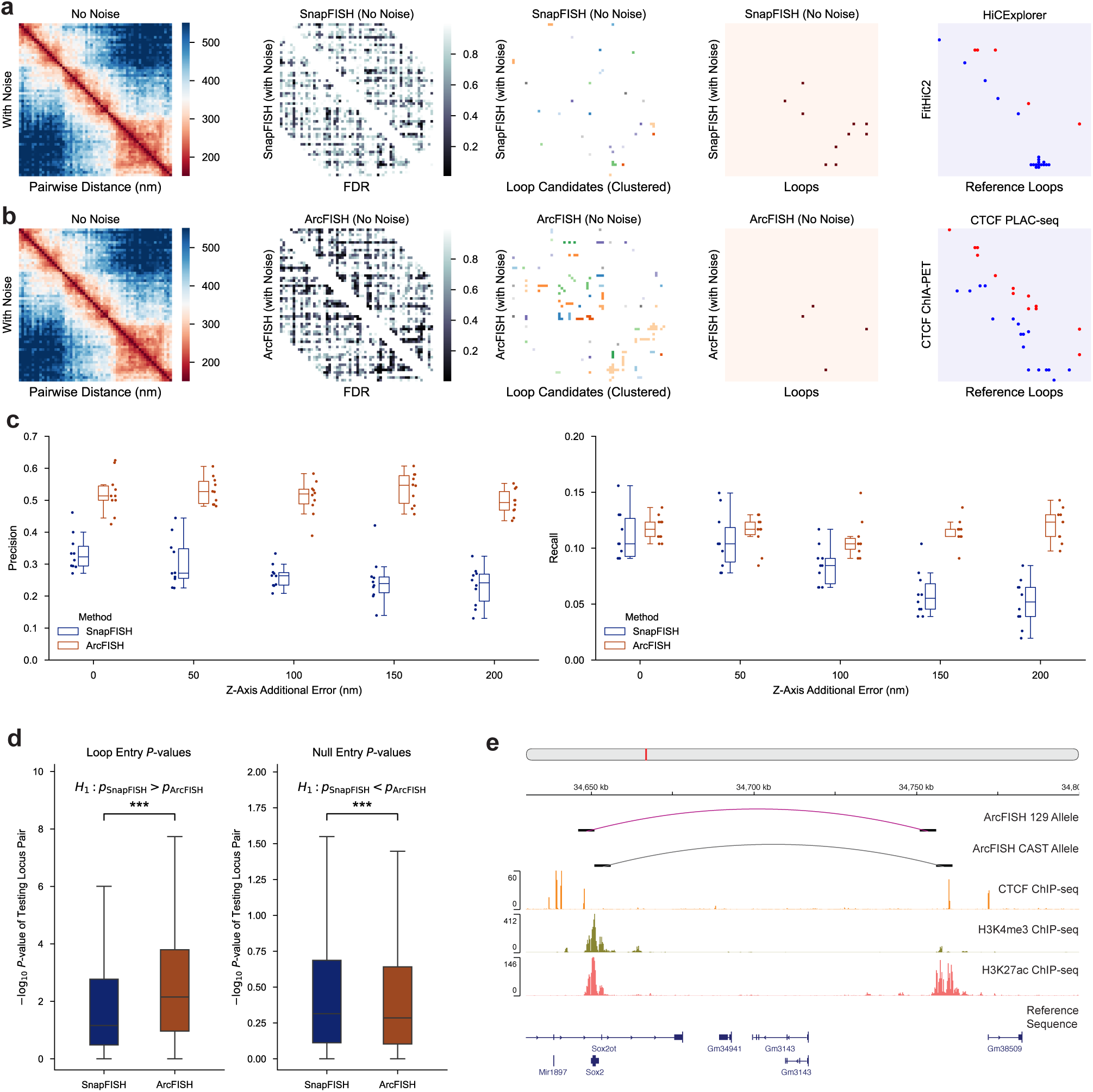
ArcFISH accurately identifies chromatin loops across different experiments. **a**, Chromosome 14 25 Kb-resolution SnapFISH loop calling results in data with uniform noise, and in data with an additional 50 nm error in Z-axis. The second heatmap is the FDR. The third heatmap is the loop candidates, colored by their clusters. The fourth heatmap is the final loops. Last heatmap is HiCExplorer and FitHiC2 identified loops from bulk Hi-C^14^. **b**, Chromosome 14 ArcFISH loop calling result. Last heatmap shows CTCF loops identified by MAPS^48^ from PLAC-seq and CTCF ChIA-PET loops. **c**, Precision and recall of SnapFISH and ArcFISH under different noise levels. We allow one 25Kb bin shift when calculating recall and precision based on the reference loops (counted as a true positive if both anchors of the identified loop are within 25Kb of one of the reference loops). N=10 samples are generated for each noise level. **d**, p-values of loop entry and non-loop entry in SnapFISH and ArcFISH. One-sided Wilcoxon signed-rank tests are performed on n=153 loop entries and n=23,814 non-loop entries. **e**, Application of ArcFISH to 5 Kb mESC chromatin tracing data. First two tracks show the position of the identified loops from the 129 allele (n=571) and the CAST allele (n=571). The next three tracks are the mESC ChIP-seq signal intensities from ENCODE^49^. The last track is the reference sequence.

We then applied both SnapFISH and ArcFISH to the original mESC data to benchmark their performance in experimental data. Excluding the X chromosome and using the same 154 loops as the working truth, SnapFISH finds 41 loops with 23 true loops (precision = 56.1%), and ArcFISH identifies 25 loops with 17 true loops (precision = 68.0%) (Figure S5). In fact, none (or 24.0% if requiring exact overlap) of loops called by ArcFISH are absent in any of the reference loop lists, while SnapFISH has 24.4% (41.5%) of loops absent in any of the reference (Figure S6a). The improvement in precision, however, might be due to the additional final p-value filtering step in ArcFISH. To make a fair comparison, we varied the p-value cutoff for both SnapFISH and ArcFISH and plot the precision-recall curve. As expected, the precision of SnapFISH increased when fewer loops are called, but its precision-recall curve is almost always bounded above by the one from ArcFISH (Figure S6b). When requiring the exact overlap between tested loops and the reference loops, the precision-recall curve from SnapFISH is always upper bounded by the one from ArcFISH, confirming that ArcFISH identifies exact chromatin loops with higher accuracy (Figure S6c). In addition, the p-values of the 154 true loops are significantly smaller in ArcFISH than SnapFISH (Wilcoxon signed-rank, p-value < 0.01), and the p-values of the remaining non-loop entries are significantly larger in ArcFISH than SnapFISH (Wilcoxon signed-rank, p-value < 0.01) (Figure 2d). These results demonstrate that loops and non-loops are more accurately distinguished by ArcFISH than SnapFISH, so the improvement is not simply due to the p-value filtering cutoff.

To evaluate the performance of ArcFISH under different resolutions and imaging protocols, we applied it to a 5Kb resolution multiplexed DNA FISH dataset in mESCs^52^. The authors imaged 42 segments around the *Sox2* promoter, and its enhancer located about 110Kb downstream. Each cell consists of the 129 allele (wild type) and the CAST allele where four CTCF binding sites were inserted between the *Sox2* promoter and enhancer. Applying ArcFISH separately to the 571 129 traces and 571 CAST traces identifies one loop for each allele as expected (Figure S7). For biological verification, we downloaded mESC ChIP-seq data of H3K27ac (active enhancer), H3K4me3 (active promoter) and CTCF^49^. Both the 129 loop and the CAST loop have their left anchor overlapped with the H3K4me3 peak at the *Sox2* gene’s promoter, and both loop anchors contain H3K27ac and CTCF peaks (Figure 2e). These together show that ArcFISH can identify biologically relevant loops at different resolutions. In addition, the p-value for the loop in the 129 allele is smaller than that in the CAST allele (4.13e-9 versus 3.26e-8), consistent with the expectation that the CAST allele-specific insertion of CTCF binding sites reduces the long-range enhancer-promoter interactions.

### ArcFISH consistently identifies domain boundaries enriched for transcription start sites and active chromatin marks

To validate our TAD calling algorithm, we re-analyzed the same DNA seqFISH+ dataset in mESCs and added 0, 50, 100, 150, and 200 nm additional error to the z-axis. We tested our ArcFISH against the insulation score based method adopted by Su et al^34^, which is the only population level TAD calling algorithm for chromatin tracing data. We refer to it as the Insulation Score method hereafter. Both Insulation Score and ArcFISH output a score for each segment based on comparing the intra- and inter-domain interactions, but ArcFISH models each axis separately. We found that ArcFISH improves the shared boundary between uniform noise data and data with additional z-axis error by 25.9% to 34.4% (Figure 3a). Even in the 50 nm error scenario, ArcFISH has a considerably higher fraction of shared boundary with the uniform error data (74.4%) compared to Insulation Score (44.8%). Since domain boundaries are enriched for regulatory elements, the performance of different TAD callers can be compared through the average count of transcription start sites (TSSs) at domain boundaries (see Methods). Indeed, ArcFISH-identified TAD boundaries have an average TSS count of about 1.9 across all noise levels, which is significantly higher than the 1.5 of Insulation Score (Wilcoxon signed-rank, p-value < 0.001) and is also higher than the average TSS count across all the imaging region (Figure 3b). Notably, ArcFISH identifies a similar number of TADs of similar sizes across different noise levels while Insulation Score is less consistent (Figure S8).

**Figure 3.**
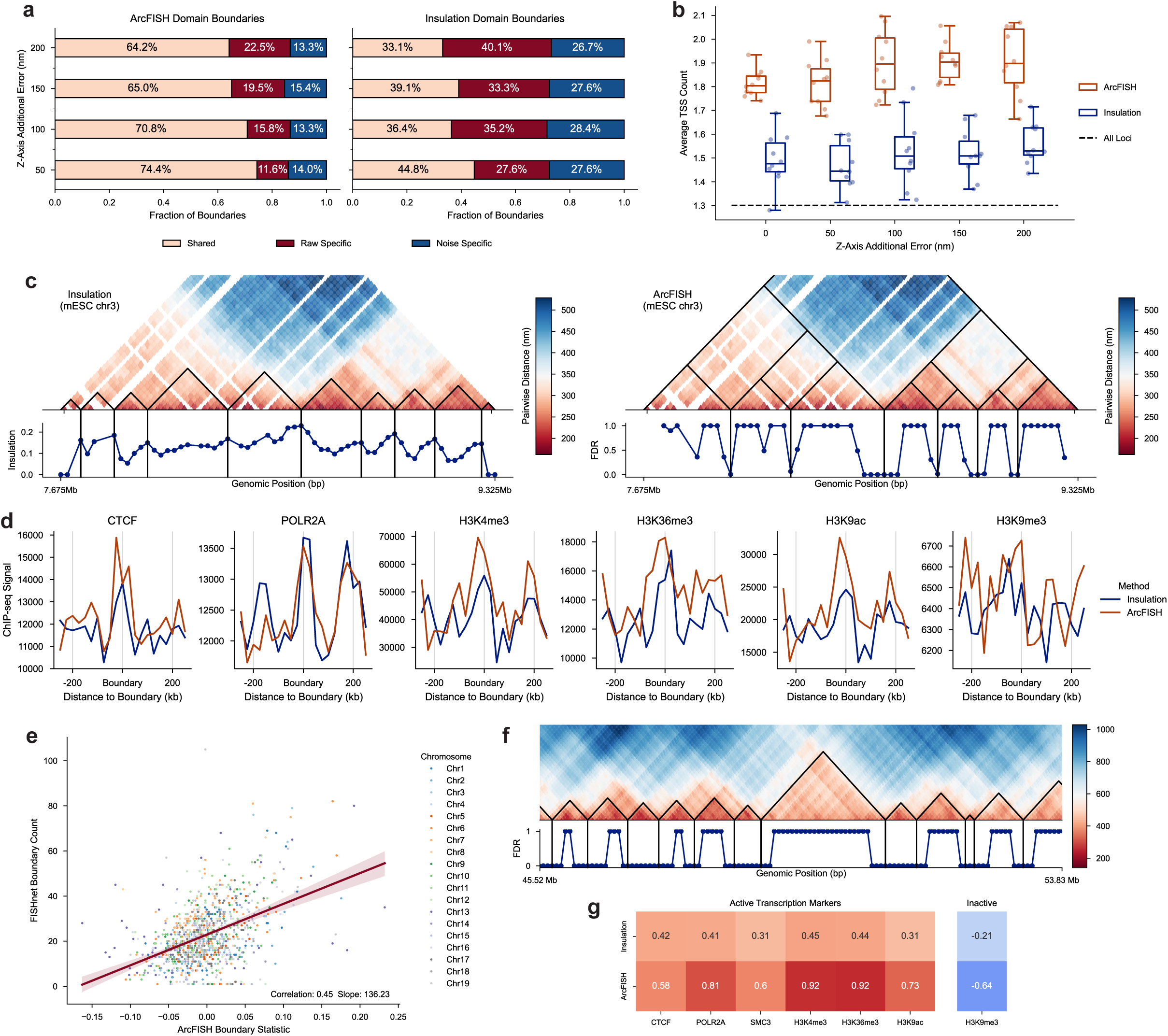
ArcFISH-identified TAD boundaries are enriched for active chromatin marks under different noise levels. **a**, Fraction of shared boundaries between data with uniform noise and data with additional Z-axis noise. “Raw specific” indicates TAD boundaries identified only in the zero-noise data, and “noise specific” indicates TAD boundaries identified only in data with additional noise. Two boundaries overlap if they are within 25Kb of each other. **b**, Average transcription starts site (TSS) count in domain boundaries detected by Insulation Score and ArcFISH. N=10 samples are generated for each noise level. Black dashed line is the average TSS count across the whole imaging region. A value higher than the black dashed line indicates TSS enrichment. **c**, DNA seqFISH+ chromosome 3 domain calling results from Insulation Score (left) and ArcFISH (right). The line plot is the insulation scores of each segment for Insulation Score and is the FDR for ArcFISH. The heatmap for ArcFISH also displays the hierarchical TADs identified by ArcFISH. ArcFISH uses an approximation of FDR to identify local peaks (see Methods). **d**, Enrichment of six epigenetic markers around Insulation Score and ArcFISH-identified TAD boundaries. The first five markers correspond to active chromatin mark while H3K9me3 is an inactive chromatin mark. **e**, Single-cell TAD boundary frequency is positively correlated with ArcFISH TAD boundary statistic. Large boundary statistics indicate potential domain boundaries in ArcFISH. **f**, Chromosome 21 domain calling results from ArcFISH. Only a ∼8 Mb region on chromosome 21 is shown. **g**, Log2 enrichment of ChIP-seq peaks in Insulation Score and ArcFISH boundaries identified from chromosome 21 chromatin tracing data.

Next, we applied ArcFISH and Insulation Score to the original mESC seqFISH+ data at 25Kb resolution. Insulation Score identifies 135 TAD boundaries, and ArcFISH identifies 105 TAD boundaries. Compared to the score defined by Insulation Score, the FDR used by ArcFISH shows clearer local minimums and is more interpretable (Figure 3c and S9). To validate the boundaries called by both methods, we downloaded the mESC ChIP-seq data^49^ of CTCF, active transcription (POLR2A), active promoters (H3K4me3), actively transcribed regions (H3K36me3), active chromatin (H3K9ac), and heterochromatin (H3K9me3) (see Methods). ArcFISH-identified TAD boundaries show a visually apparent enrichment for CTCF and all active transcription markers (POLR2A, H3K4me3, H3K36me3, and H3K9ac) despite the relatively small number of TADs (Figure 3d). Moreover, the peaks of H3K4me3 and H3K9ac are substantially higher than the ones from Insulation Score, showing that ArcFISH identifies more biologically relevant TAD boundaries. For the inactive transcription marker H3K9me3, it is hard to claim it is depleted or enriched due to large variation (Figure 3d). One reason is the imaging region only has 47 segments with H3K9me3 marker while other markers overlap at least hundreds of segments.

The limited segment number likely leads to the false enrichment. ArcFISH also achieved higher ChIP-seq peaks enrichment as expected (Figure S10). To further benchmark the ArcFISH-identified TADs, we compared them with the FISHnet-identified TAD-like structure in single cells (see Methods). Indeed, TAD boundaries with larger test statistics in ArcFISH tend to appear more frequently in single cells (Pearson correlation = 0.45, p-value < 0.001), irrespective of chromosomes (Figure 3e and S11). In summary, through multiple data sources and methods, we validate ArcFISH-identified TAD boundaries are biologically meaningful.

In addition, we benchmarked ArcFISH on human chromosome 21 multiplexed chromatin tracing data from Su et al and evaluated the effect of sample size on TAD calling. Su et al imaged 10.4-46.7 Mb of chromosome 21 with 651 probes at 50Kb resolution, generating a total of 7,591 traces. We applied Insulation Score and ArcFISH and identified 53 and 49 TAD boundaries, respectively (Figure 3f and S12). ArcFISH-identified TAD boundaries achieved a higher enrichment with CTCF, SMC3, and active histone marks POLR2A, H3K4me3, H3K36me3, and H3K9ac compared to Insulation Score (Figure 3g). For inactive chromatin marks, H3K9me3 is depleted in ArcFISH-identified TAD boundaries with a log2 fold change of -0.56, more than twice smaller than the -0.21 from Insulation Score. This shows the previous enrichment of H3K9me3 is likely due to insufficient samples and confirms that ArcFISH consistently outperforms Insulation Score in identifying transcription correlated domain boundaries. To test the influence of the number of available chromatin traces, we selected 100, 200, 400, 800, 1,600, 3,200, and 6,400 traces, applied ArcFISH on 10 random samples for each number of traces, and calculated the Pearson correlation coefficients between the FDR profile from the sample and the one from the full data. The correlation is close to 0.9 when there are only 200 traces and becomes higher as the sample size increases (Figure S13a). FDRs converged to 0 or 1 as sample size increases, showing the statistical test can clearly separate boundaries and non-boundaries (Figure S13b). The recall and precision, evaluated by treating the boundaries from the full data as the working truth, confirm that 200 to 400 traces are sufficient for reliable TAD calling (Figure S13a). Indeed, the correlations with ChIP-seq markers remain high under different numbers of input traces (Figure S13c). Therefore, ArcFISH is capable of accurately identifying TAD boundaries with a limited number of traces.

### ArcFISH enables accurate A/B compartment identification by adaptive weighting

We used chromosome 2 (chr2) chromatin tracing data from Su et al to benchmark the A/B compartment calling performance of ArcFISH. The first dataset contains 4,848 traces spanning the p-arm of chromosome 2. Each trace contains 357 imaging segments at 250Kb resolution. The second dataset contains 3,029 traces covering the entire chromosome 2 by 935 imaging segments, again at 250Kb resolution. Both datasets exhibit larger measurement uncertainties in the x and y axes than in the z-axis (Figure S14a and S14b). In the original paper, the authors converted the pairwise distance matrices into a pseudo-count matrix and assigned A/B compartments by the sign of the first principal component (PC) from the correlation matrix. We first applied ArcFISH to the dataset of chr2 p-arm and compared the compartment assignments with those from PC1 (Figure S15a). Out of the 357 segments, only 27 segments (7.6%) are assigned differently by the two methods (Figure 4a). We then validated the A/B compartments using histone modifications from mESC ChIP-seq data. As expected, both methods associate A compartments with active chromatin and B compartments with inactive chromatin, and the log2 fold changes of ChIP-seq peaks are similar (Figure 4b and S15b). The results show that both methods can identify A/B compartments well with a relatively large sample size (thousands of chromatin traces).

**Figure 4.**
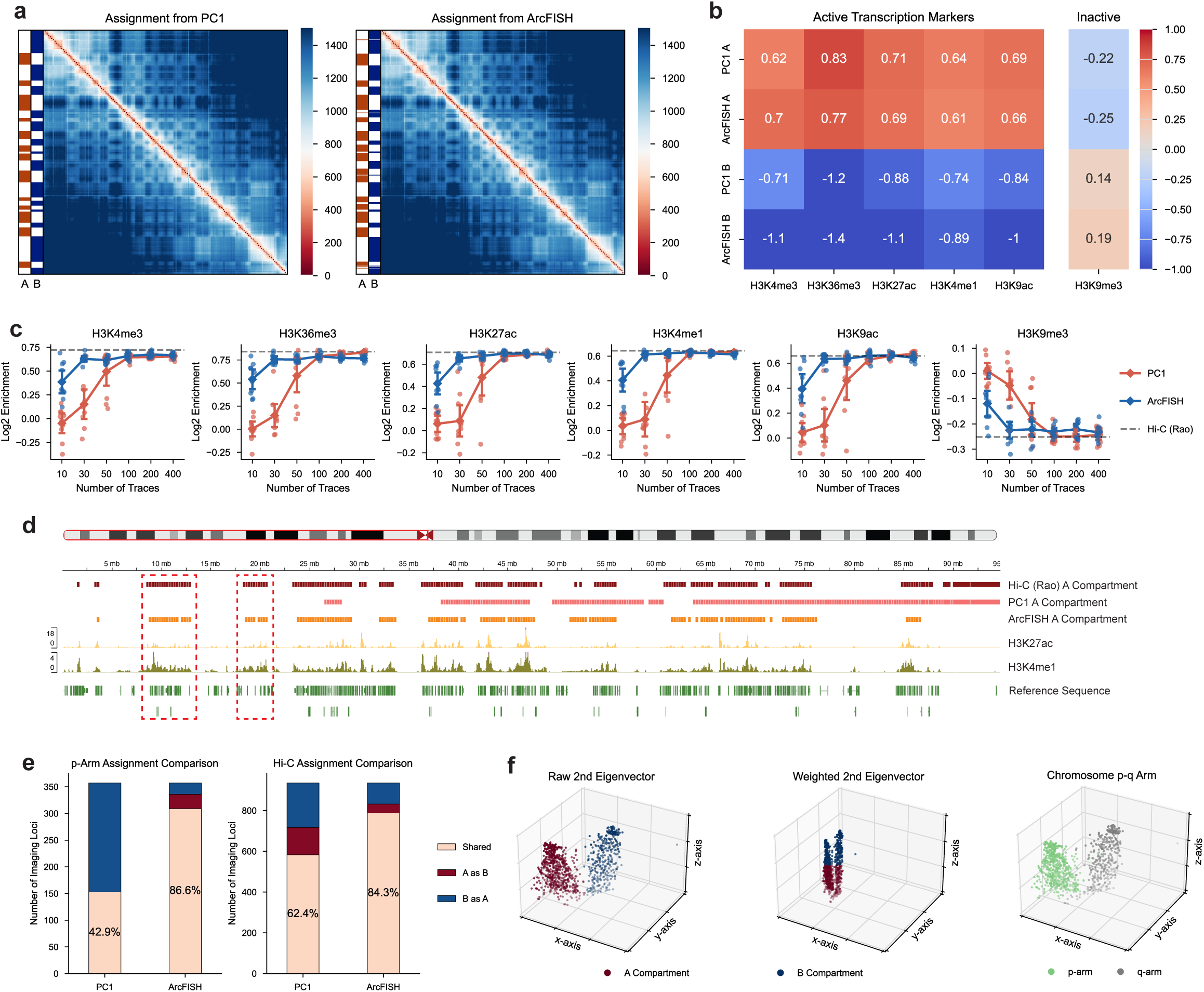
ArcFISH identifies A/B compartments using fewer traces and the entire chromosome through adaptive weighting. **a**, A/B compartment assignments of chromosome 2 p-arm by PC1 and ArcFISH. Heatmaps show the median pairwise distance in nm. **b**, Enrichment of histone marks in A and B compartments of chromosome 2 p-arm. The enrichment is evaluated by the log2 fold change of ChIP-seq peaks. **c**, Log2 enrichment of histone marks in A compartment across different number of traces. Error bars represent the standard error from n=10 replicates for each number of traces. Grey dashed line is the log2 enrichment in bulk Hi-C data. **d**, A compartment assignment of chromosome 2 p-arm using 30 traces. The first three tracks are the A compartment assignments from bulk Hi-C, PC1, and ArcFISH. The next two tracks are histone marks enriched in A compartments, and the last track is the reference sequence. Red boxes highlight regions with active chromatin markers that PC1 fails to capture but captured by bulk Hi-C and ArcFISH. **e**, Shared compartment assignments between entire chromosome 2 and p-arm alone or bulk Hi-C. Left bar plot shows the agreement of PC1 and ArcFISH between entire chromosome 2 and p-arm alone. Right bar plot shows the agreement of PC1 and ArcFISH between entire chromosome 2 and bulk Hi-C. **f**, Second eigenvector from each axis. First scatter plot is the A/B compartment assignment based on K-means from the raw second eigenvectors. Second scatter plot is the A/B compartment assignment based on K-means from the weighted second eigenvectors. Third plot colors each segment by p-q arms.

We next compared the A/B compartment assignment results with smaller numbers of traces. We randomly selected 10 samples for 10, 30, 50, 100, 200, and 400 traces and compared the histone mark enrichment across all replicates. Both methods achieved similar performance (measured by the levels of histone mark enrichments in compartment A) when the number of traces was more than 100, reaching the level of accuracy comparable to deeply sequenced bulk Hi-C data^10^. Notably, the two methods differ in their performance with fewer traces (Figure 4c, S16a and S17a). For instance, ArcFISH has an average log2 fold change of 0.757 for H3K36me3 (active transcribed regions) with 30 traces while the log2 fold change is only 0.147 in PC1 (Mann-Whitney, p-value < 0.001). As a reference, the A/B compartment annotation from the deeply sequenced bulk Hi-C data has a log2 fold change of 0.841, which is comparable to the value from ArcFISH. The same pattern occurs in other histone marks with statistically significant differences (p-value < 0.05 for all markers). When looking into specific replicates, ArcFISH A/B compartment assignments have more alternations between A and B as observed in Hi-C annotations, and ArcFISH compartment A captures more H3K27ac and H3K4me1 peaks compared to PC1 (Figure 4d). These results suggest ArcFISH can accurately identify A/B compartments with a small number of chromatin traces.

To further demonstrate the necessity and benefits of axis weighting, we applied both PC1 and ArcFISH to the second dataset, which consists of 935 imaged segments on the entire chromosome 2. Instead of computing PC1 on the p and the q arms separately as was done in the original paper, we re-analyzed the entire dataset to investigate whether the methods are capable of consistently assigning compartments, irrespective of the potential confounding factor of the two chromosome arms. As expected, PC1 assigns A/B compartments by p-q arms, with only 42.9% common assignments compared to applying PC1 to the p arm alone and 62.4% agreement with bulk Hi-C (Figure S16b and 4e). In contrast, ArcFISH is robust to the presence of both the p arm and the q arm. It has 86.6% common assignments compared to applying ArcFISH to the p arm alone and 84.3% shared assignments with bulk Hi-C (Figure S16b and Figure 4e). In addition, ArcFISH achieved higher enrichments of ChIP-seq histone marks compared to PC1 (Figure S17b).

To understand why ArcFISH is more robust to two chromosome arms, we plotted the second eigenvector from each axis, which ArcFISH uses to perform K-means clustering and output final A/B compartment assignments. We also plotted the p arm and the q arm in the 3D space for visual comparison. As shown in Figure 4f, the p arm and the q arm clearly separate in the x and the y axes, and K-means clusters A/B compartments as p-q arms in the raw 3D space as expected. However, when we applied the weights estimated by ArcFISH to different axes, the z-axis is stretched more than the other two axes, leading to a larger clustering weight. Therefore, K-means clustering can distinguish A/B compartments based on the more accurate information from the z-axis. This is also reflected by the relative enrichment of histone markers (Figure S17c). In sum, ArcFISH effectively leverages information by adaptively assigning axis weights, enabling accurate and robust A/B compartment identification.

### ArcFISH identifies cell-type-specific TADs and chromatin loops

As an application of ArcFISH, we compared the TADs and chromatin loops identified from different cell types in the mouse cerebral cortex. Using the same 25Kb segments as in mESCs, Takei et al imaged ∼1.5 Mb regions on each chromosome for 9 cell types across three biological replicates. Although thousands of single cells are imaged, most cell types have fewer than 250 chromatin traces available. However, ArcFISH still identifies valid TAD boundaries, and the hierarchical TAD structure varies among cell types. For example, Figure 5a shows a TAD reorganization from Pvalb to Astro. To further investigate whether TAD boundaries are cell-type-specific at 25Kb resolution, we applied ArcFISH to each of the three biological replicates and performed dimension reduction based on the FDR profiles. In the t-SNE plot, biological replicates from the same cell type cluster together (Figure 5b). Notably, for Astro, Endo, and Pvalb, each biological replicate contains only ∼50 to ∼120 chromatin traces, indicating the high sensitivity of ArcFISH. We also compared the chromatin loop calling results for different cell types. As expected, the number of loops identified depends on the number of available traces (Figure S18). To make a fair comparison between cell types, we visualize the top 1%, 3%, and 5% FDR thresholds instead of setting a uniform FDR cutoff. Like TADs, chromatin loops are also cell-type-specific and reflect the differences observed in median pairwise distance maps (Figure 5c).

**Figure 5.**
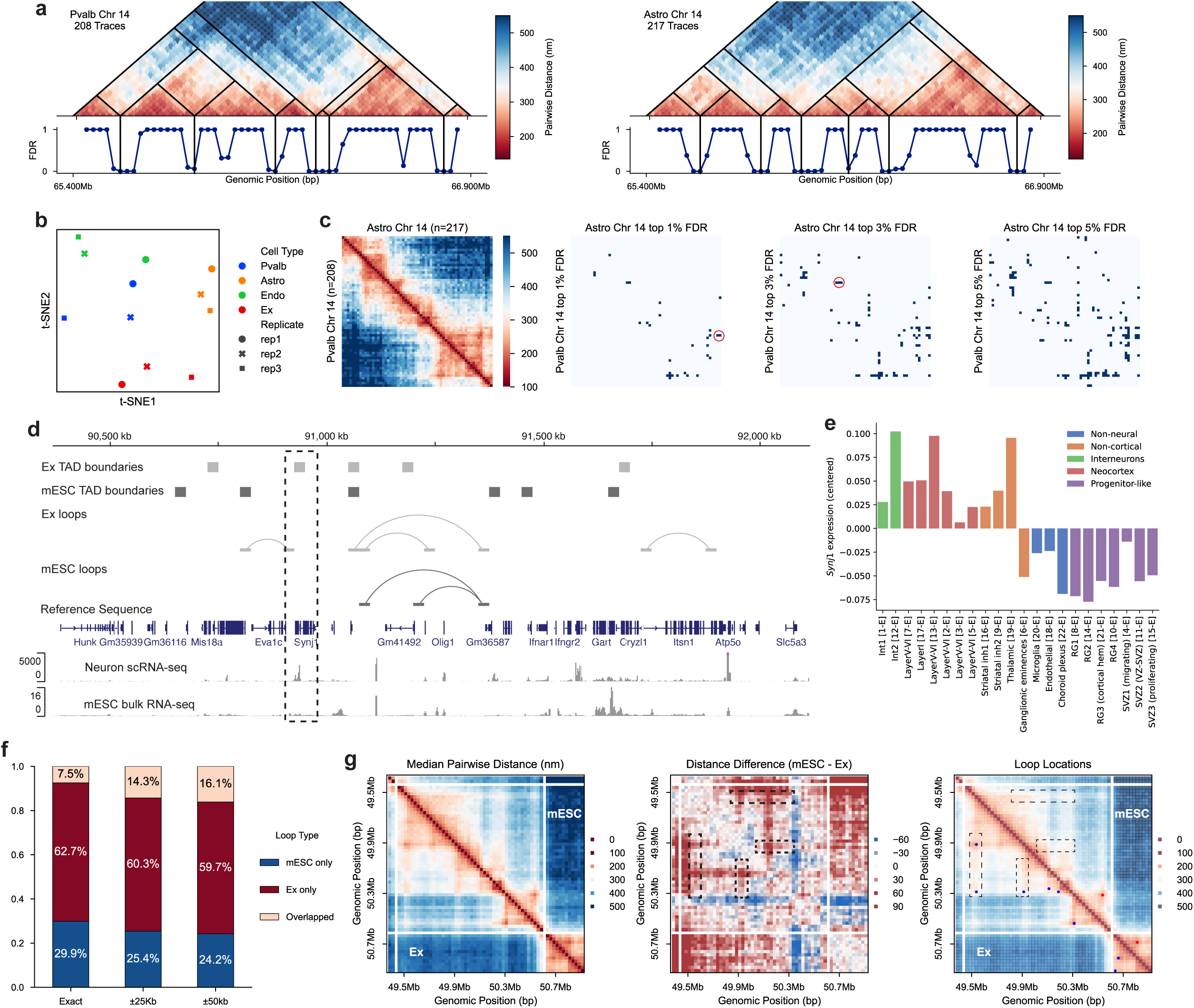
ArcFISH reveals cell-type-specific TADs and chromatin loops. **a**, DNA seqFISH+ chromosome 14 TAD calling results for Pvalb and Astro. The hierarchical TAD structure shows change in preferential TAD arrangement. **b**, t-SNE plot of TADs from each biological replicate. Each dot represents a biological replicate from a cell type. t-SNE is performed on the TAD FDR of chromosome 1 to 19. **c**, Different cell types have distinct chromatin loops at the same significance level. The upper left heatmap is the median pairwise distance matrices for Astro and Pvalb. The remaining are the chromatin loop FDRs binarized at the same proportion. **d**, *Synj1* lies within a differential TAD boundary and a differential chromatin loop. The single-cell RNA-seq of mouse neurons is provided by the integrative genome browser^71^, and the bulk RNA-seq of mouse embryonic stem cells is obtained from GEO with accession number GSE75804. **e**, Relative expression of *Synj1* in different cell types. The y-axis is the average fold change of expression relative to housekeeping genes, centered to zero. **f**, The proportion of shared chromatin loops between mouse embryonic stem cells and excitatory neurons. Exact means exact overlap; ±25Kb means allowing one 25Kb bin shift; and ±50Kb means allowing 50Kb shift. **g**, Excitatory neurons have more loop extrusion activities than mESCs. Chromosome 6 imaging region is shown. The middle heatmap is the difference of median pairwise distance matrix. Black boxes highlight chromatin stripes present in excitatory neurons but not in mESCs.

To understand how 3D genome structure changes as cells differentiate, we compared TADs in mESCs and mouse excitatory neurons. At the same sample size, there was no significant difference in the number of TAD boundaries (101 versus 115, Fisher exact p-value = 0.353), TAD size (Mann-Whitney, p-value = 0.508), and transcription start sites (TSS) count at TAD boundaries (Fisher exact p-value = 0.778), showing that the two cell types have similar TAD distribution at the 25Kb resolution (Figure S19a and S19b). We next analyzed the histone mark enrichment in cell-type-specific and shared TAD boundaries. Although no statistical significance was detected due to the relatively small imaging region, we observed a general trend that active histone marks are more enriched in shared TAD boundaries, particularly for marks associated with active promoters (Figure S19c).

We also investigated whether genes at differential TAD boundaries are expressed in a cell-type-specific manner. As an illustrative example, we focused on the *Synj1* gene on chromosome 16, which lies within the TAD boundary of excitatory neurons but not mESCs (Figure 5d and S20). *Synj1* encodes a polyphosphoinositide phosphatase enriched in neurons, and heterozygous deletion of this gene leads to Parkinson’s disease-like pathologies in mice^53,54^. To evaluate the expression of *Synj1*, we checked two published scRNA-seq datasets of excitatory neurons^55^ and mESCs^56^. Since the two datasets originated from different batches, typical differential gene analysis pipelines do not apply. Instead, we identified a list of housekeeping genes^57^ and calculated the fold change of *Synj1* expression relative to the average expression of housekeeping genes (see Methods). The mean log fold change in excitatory neurons across all cells is -0.053 while the one in mESCs is -0.320, and the average counts per million (CPM) of *Synj1* is 55.68 in excitatory neurons and 26.04 in mESCs, showing that *Synj1* is more highly expressed in excitatory neurons than in mESCs. Moreover, *Synj1* is differentially expressed in excitatory neurons and glial cells though the TAD boundary also occurs in glial cells (Figure 5e). In short, differential boundaries found by ArcFISH mark potential differentially expressed genes.

We next compared the chromatin loops in mESCs and mouse excitatory neurons. In contrast to TADs, at the same sample size, we identified significantly more chromatin loops in excitatory neurons than in mESCs (47 versus 25, Fisher exact p-value = 0.006), and most loops are differential (Figure 5f). This is consistent with previous observations from Hi-C data that chromatin loops show more cell type variability than TADs^10^. Visually, the pairwise distance matrix from excitatory neurons has more apparent chromatin stripes^58^ than the one from embryonic stem cells, and the additional loops identified by ArcFISH typically correspond to the end of such stripes (Figure 5g). Since stripes indicate potential stalled SMC complexes, this might imply that excitatory neurons have more loop extrusion activities in the given imaging regions.

## Discussion

In this paper, we describe a new computational method ArcFISH, which can identify chromatin loops, TADs, and A/B compartments from chromatin tracing data. By explicitly modeling each axis separately and adaptively weighting axis-specific measurement errors, ArcFISH can accurately and robustly identify 3D genome features compared to existing methods, which are primarily adapted from computational tools developed for Hi-C data. We benchmark ArcFISH by applying it to multiple datasets across different cell types in both mice and humans, and multiple imaging protocols. For chromatin loops, we systematically evaluated recall and precision across various noise levels. For TADs, we used histone ChIP-seq data and transcription start sites to justify the biological relevance of ArcFISH-identified TAD boundaries. The performance of ArcFISH in identifying TAD boundaries across various sample sizes and noise levels is highly consistent, overcoming limitations of the Insulation Score. For A/B compartments, we benchmarked the results using histone modification ChIP-seq data and found that using as few as 30 chromatin traces, ArcFISH identifies A/B compartments consistent with that obtained from bulk Hi-C data. Finally, as an application, we applied ArcFISH to a mouse brain imaging dataset and identified cell-type-specific TADs and chromatin loops, revealing that excitatory neurons have more loop extrusion activity compared to mouse embryonic stem cells.

The main innovation in ArcFISH lies in modeling axis-specific measurement errors and developing a unified statistical framework to leverage information from the more accurate axes. While ArcFISH adopts the same testing region as Insulation Score for TAD calling and similar PCA rationale for A/B compartment calling, in principle, any computational methods developed for Hi-C data can fit into this general framework to efficiently process chromatin tracing data. When analyzing new chromatin tracing datasets, our findings suggest even one or two axes contain substantial measurement error, we can still extract meaningful information if there is one accurately measured axis. This implies that to improve the resolution or the quality of chromatin tracing data, it is not necessary to simultaneously scaling up all three axes. Instead, focusing on one or two axis is sufficient, which is also much easier in practice.

ArcFISH can be further improved in the following directions. First, while the existing software can efficiently store and analyze chromatin-tracing datasets containing thousands of imaging segments and tens of thousands of single cells, more recent experiments can capture hundreds of thousands of segments^25^. To enhance scalability, future versions of ArcFISH could employ Dask arrays throughout downstream processing—not just during preprocessing and normalization Second, computational tools for downstream tasks are needed. After defining population level 3D genome features, we may focus on single-cell level variability. We can also integrate with other data modalities such as gene expression (RNA MERFISH, RNA seqFISH+), or cell surface protein marks (sequential immunofluorescence), which are usually provided together with chromatin tracing data, to investigate the relation between 3D genome and gene regulation.

## Methods

### Chromatin trace model and motivation of ArcFISH

Let (***x, y, z***) ∈ ℝ*^p^*^X3^ denote the observed 3D coordinates of a chromatin trace of length *p*. We first center the chromatin trace by subtracting each axis by its mean. The observed 3D location consists of two parts, the true 3D location 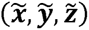 and the measurement error (**ε***_X_,***ε***_y_,***ε***_z_)*. We further assume that both parts follow Gaussian distribution, that is, 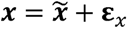 with 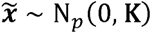 and 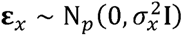 and similarly for the other two axes. Though the distribution of 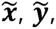 and 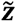 are the same since they are different views of the same chromatin, the error terms **ε***_X_,***ε***_y_,* and **ε***_z_* have different variance 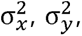 and 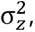 thus capturing the distinct measurement errors observed in real chromatin tracing data.

Existing approaches^22,34–36^ to identify 3D genome features operate on the pairwise distance matrix D with 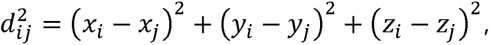 an unweighted sum of squared pairwise difference matrices. Since pairwise distance contains information regarding 3D genome features, each axis pairwise difference matrix also characterizes loops, TADs, and A/B compartments. Therefore, it is feasible to first analyze each axis separately and combine the results with appropriate weights. In contrast, methods based on the pairwise distance treat each axis equally as the summation is unweighted, thus lacking the ability to prioritize more accurate axes. We describe the details of each step in ArcFISH below.

### Axis weight estimation

As explained in the main text, an ideal choice of weights is the inverse of measurement errors, but 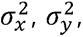 and 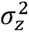 are not identifiable without further assumptions (Supplementary Methods Section 1). Like Hi-C contact matrices, where some methods^59–61^ only use the top few eigenvectors and treat the rest as noise, we assume that the squared pairwise distance matrix is inherently low rank as well. With the additional assumption that its rank is smaller than *p*/2 - 2, the measurement errors become identifiable Supplementary Methods Section 1). In fact, the model possesses a spiked covariance structure^62^. To further accommodate the high-dimensional setting, where the number of imaged segments is comparable or larger than the number of available traces, we adopt the estimator described in the sparse PCA paper^63^.

Specifically, for each axis, we calculated the sample covariance matrix and used the median of its diagonal elements as the estimated measurement errors. Then we set the axis weight *w_X_*, *w_y_*, and *w_z_* as the inverse of the estimated measurement error and normalized them so that they sum to one.

### Filtering and normalization

To denote all *n* traces in a chromatin tracing dataset, we add a subscript *i* so that (***x_i_, y_i_, z_i_***) ∈ ℝ*^p^*^X3^ is the 3D coordinates of the *i*th chromatin. We first describe how abnormal values in pairwise difference matrices (values of the form *x_ij_* - *x_ik_*, where *j* and *k* are imaged segment indices, and similarly for the other two axes) are filtered. Essentially, pairwise difference values are grouped by 1D genomic distance, and within each 1D distance stratum, values with a Z-score larger than 4 or smaller than -4 are filtered out. When implementing the filtering step, however, we rely on an intermediate raw axis variance matrix to represent all pairwise difference values, which help speed up computation. We also use a lowess curve when estimating stratum standard deviations since some stratum contains only few valid values.

Specifically, for each axis, we first compute the raw *p* x *p* axis variance matrix whose *jk*th entry is the sample variance of *x_ij_* - *x_ik_*, except that all the mean operations are replaced by the median to improve robustness. A lowess curve is then fitted to the raw axis variance matrix against the 1D genomic distance to estimate the expected variance at each fixed genomic distance. Then *x_ij_* - *x_ik_*’s farther than four expected standard deviations away are treated as outliers and filtered out. Second, based on the filtered pairwise difference matrices, we compute filtered *p* x *p* axis variance matrix whose *jk*th entry is the sample variance of *x_ij_* - *x_ik_*. A lowess curve is then fitted to the filtered axis variance matrix against 1D genomic distance. Finally, the filtered axis variance is divided by the predicted axis variance to remove 1D genomic distance bias. This results in a *p* x *p* normalized axis variance matrix for x, y, and z axes. We made some modifications in the following steps (Supplementary Methods Section 4) so that all operations can be performed using only the normalized variance matrix, without accessing the raw pairwise difference matrices (*n* x *p* x *p* tensors, scales linearly in the number of traces and quadratically with the number of segments).

### Chromatin loop calling

ArcFISH employs a local background model to identify chromatin loops, similar to HiCCUPS^14^, HiCExplorer^50^, and SnapFISH^36^. For each segment pair *(j, k)*, its local background contains all segment pairs with a 1D genomic distance of 25 Kb to 50 Kb from the tested segment pair (loop segments, i.e., the *j*th and *k*th segments) (Figure S2). If the tested segment pair is a loop, then the 3D distance *d_jk_* should be significantly smaller than the 3D distance *d_j’k’_* in the local background, where *j’k’* ∈ *B_jk_* and *B_jk_* is the index set of the local background. If *B_jk_* contains fewer than 3 entries, the corresponding segment pair is not tested due to lack of samples. Since *d_jk_* is a sum of the squared axis difference as described previously, we can test each axis separately. Specifically, for each axis, we pool all the background entries together and calculate a background sample variance defined as:

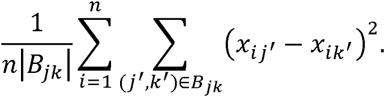

Similarly, a sample variance is computed for the tested segment pair. Due to missing values, the denominator is adjusted based on the number of available pairwise differences. Under the null hypothesis that the tested segment pair and the segment pairs in the local neighborhood region have the same variance, the ratio between the tested segment pair sample variance and the background sample variance follows a F-distribution, whose degrees of freedom equal to the denominators in the sample variances (Supplementary Methods Section 2). The F-test results in a p-value for each axis, *p_X_*, *p_y_*, and *p_z_*. We then use aggregated Cauchy test^43^ to combine the axis-specific p-values:

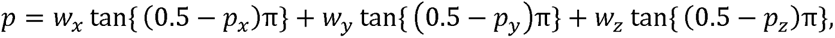

where *w_X_*, *w_y_*, and *w_z_* are the axis weights estimated in the first step.

After computing p-values for all segment pairs, we convert them to false discovery rates (FDR) to adjust for multiple testing and define a segment pair as a loop candidate if its FDR is smaller than 0.1. Loop candidates within 50 Kb of each other are grouped together, and the segment pair with the smallest p-value is designated as the loop summit, while other candidates are filtered out to control false positives. Since the loop-calling results will be compared to Hi-C loop calling results, which are subject to a higher multiple testing burden as they are applied to the whole genome, we include an additional final filtering step based on p-values. Specifically, we retain only loop summits with p-values smaller than 1e−5, which is comparable to the loop p-values generated by FitHiC2^51^ when applied to mouse embryonic stem cell Hi-C data from Bonev et al^14^. For seqFISH+ and Huang et al^52^ data, we further filtered out cluster summits with a population-level contact frequency less than 1/3 and singleton summits (50 Kb neighborhood only contains one summit) with a population-level contact frequency less than 1/2 for a fair comparison with SnapFISH^36^.

### Domain calling

For TAD calling, ArcFISH outputs a p-value for each imaged segment indicating whether the segment is a TAD boundary or not. For each segment, we adopt a fixed length window preceding and after the segment. If the segment is at the two ends of the imaged region so that more than half of the window is unavailable, the segment is excluded for domain boundary calling. For the rest of the imaged genomic segments, like Insulation Score^42^, we compared the inter-domain and intra-domain 3D distance. If the testing segment is a TAD boundary, the inter-domain 3D distance should be significantly larger than the intra-domain 3D distance since segments upstream and downstream of the tested segment interact more frequently within their respective domains than across. Using the same distance-variance relationship, we tested the variance of each axis separately. For each axis, we pooled the inter-domain pairwise difference together and calculated an inter-domain sample variance. Similarly, an intra-domain sample variance is obtained for each axis. Then the ratio between the two variances follows a F-distribution under the null hypothesis that the tested segment is not a TAD boundary (Supplementary Methods Section 3). As in chromatin loop calling, the F-test outputs a p-value for each axis, and we combined the axis-specific p-values into a single p-value by the aggregated Cauchy test. The p-values are converted to FDRs to adjust for multiple testing, and segments with FDRs smaller than 0.1 are defined as the candidate TAD boundaries.

Since segments near a TAD boundary typically has small FDR values, we only kept local minimum of TADs, using the `find_peaks` function from `scipy.signal`^64^ on the FDR profile (a length *p* vector). However, we observed that the FDRs become all zeros and ones as the number of traces increases to few thousands, leading to numerical instability of peak calling.

Thus, instead of using the raw FDR profile, we calculated approximated p-values based on an F-distribution with both degree of freedoms equal to one (instead of the sample sizes) and performed peak calling on the resulting approximated FDRs. Segments with raw FDRs smaller than 0.1 and marked as peaks are kept as the final TAD boundaries.

We also include an algorithm to define hierarchical TADs (Figure S3). First, the region between each consecutive TAD boundary pair is included in the final TAD list with level equal to zero. Next, we sequentially removed boundaries ranked by their aggregated Cauchy test statistics, with significance level from low to high, and added the region between the immediate boundary preceding and after the removed boundary to the TAD list with level increased by one each time (Figure S3). This procedure is repeated until all TAD boundaries are removed. The final TAD list, along with the assigned levels, provides a hierarchical structure that can be visualized or filtered accordingly.

### Compartment calling

We model the A/B compartment with a two-block stochastic block model (SBM)^65^. Briefly, we treat each segment as a node with an unknown membership—A or B—and segments with the same membership tend to interact with each other more frequently. To infer the node membership, we used spectral clustering, a widely adopted method in SBM literature with well-established statistical properties^66,67^. However, instead of applying clustering to the top two eigenvectors from a single similarity matrix, as in the standard spectral clustering, we performed eigenvalue decomposition separately for each axis and clustered in the resulting joint 3D feature space. Specifically, we extracted the second eigenvectors from the median-normalized pairwise difference matrices of each axis to construct a three-dimensional feature space (Supplementary Methods Section 5). We then scaled each feature by the corresponding axis-specific weight and applied the K-Means clustering to the weighted feature space. The two resulting clusters from K-Means are the final A/B compartment assignments. The additional weighting step ensures that the eigenvectors from more accurate axes are upweighted in the final clustering step, leading to an improved A/B compartment assignment.

After clustering the segments into two groups, it remains ambiguous which group corresponds to the A compartment and which group corresponds to the B compartment. If the gene annotation is available in the UCSC genome browser^68^, ArcFISH fetches the corresponding sequencing data and calculates the number of TSSs overlapped with the two groups. The group with a higher TSS overlap is assigned as the A compartment. ArcFISH also allows calling A/B compartment separately for the p and the q arms when a reference genome is provided (set `centromere=True`). If a reference genome is not provided or it cannot be found at the genome browser, the group with lower within-group axis variance is assigned as B compartment as heterochromatin is not transcriptionally active and thus condensed. However, it is highly recommended to use TSS to assign A/B compartments as it is much more accurate.

### Chromatin tracing data processing

Due to allele misalignment, seqFISH+ data from Takei et al^23,24^ can contain multiple 3D coordinates for the same imaged segment on the same allele. In our analysis, only the first 3D coordinate is kept while the remaining duplicated rows are removed.

Multiplexed DNA FISH data from Huang et al^52^ has an additional imaged segment (the 25^th^ segment) on its CAST allele as it has an CTCF insertion. However, we found that in the data deposited at 4DN data portal, the 129 allele also has 3D coordinates of the 25^th^ segment due to potential misalignment, and the CAST allele has abnormally high detection efficiency of the 25^th^ segment. To avoid potential technical confounding, we removed the 25^th^ segment in both alleles.

### Real data-based simulation of noisy chromatin tracing data

To evaluate the effect of different levels of noise heterogeneity, we first simulated a uniform noise chromatin tracing data from Takei et al^23^. Specifically, we only kept the x-axis coordinates and sampled three *p*-dimensional vectors from the x-axis. These three vectors together then serve as a new simulated chromatin trace (a *3x p* matrix). As the new x, y, and z coordinates in the simulated data all come from the x-axis in the original data, they have the same level of measurement uncertainty. We then added different levels of Gaussian noises to the z-axis of the simulated data and used it to benchmark different methods. Regarding the choice of additional noise, we selected 0, 50, 100, 150, and 200 nm as the standard deviation of Gaussian noises. This is supported by the following evidence: 1) the median displacement distance (distance between fluorescence signals of the same segment in different imaging rounds) is typically around 30 nm to 70 nm^22,27,34,45^; 2) displacement errors are typically right-skewed^27^, so the mean is higher than the median; 3) and there are other sources of measurement errors such as correction of drift and chromatin aberrations.

### Working truth for chromatin loop calling

Given the broad spectrum of biological technologies and computational tools to identify chromatin loops^69^, we collected and curated six representative loop lists instead of relying on a single reference set. We selected two loop lists for each of the three major technologies: PLAC-seq, ChIA-PET, and bulk Hi-C. For PLAC-seq, MAPS-identified loops targeting CTCF and H3K4me3 are downloaded from Juric et al^48^ and filtered to keep only loop summits and singletons. This resulted in 158 CTCF loops and 238 H3K4me3 loops. For ChIA-PET, CTCF and POLR2A loops are downloaded from ENCODE with IDs ENCFF112DXO (CTCF) and ENCFF877XWN (POLR2A). We kept loops with counts greater than five to avoid ChIA-PET loops dominating the reference loop lists, leading to 243 CTCF loops and 297 POLR2A loops. For Hi-C, we downloaded 10Kb resolution Hi-C data from the 4DN data portal with ID 4DNFI4OUMWZ8. We ran HiCExplorer using the recommended parameters `--windowSize 10 --peakWidth 6 --pValuePreselection 0.05 --pValue 0.05` and FitHiC2 using one spline pass, FDR smaller than 0.01, and `merge-filter` on output loops. This gave a total of 55 HiCExplorer loops and 213 FitHiC2 loops. We chose HiCExplorer and FitHiC2 since they represent the local background model and global background model typically used in Hi-C loop calling.

The six loop lists are then mapped back to the imaged segments used in Takei et al^23^. For each loop list, each loop is compared with all possible combinations of imaged segment pair and checked overlap. If both loop anchors overlap with the imaged segment pair, then the segment pair is marked as presented in the reference loop list once. After checking all six loop lists, segment pairs presented in the reference lists at least twice are included in the final loop list. This resulted in a total of 154 final loops.

### Overlap between TSSs and TAD boundaries

The TSS annotation for mm10 is downloaded from the UCSC genome browser^68^. For each 25 Kb imaged bin in Takei et al^23^, we counted how many TSSs overlapped with the imaged bin. We then calculated the mean TSS count over all TAD boundary bins.

### Overlap between ChIP-seq and TAD boundaries

ChIP-seq data to validate mESC TADs is downloaded from ENCODE^49^ with the following IDs: H3K4me3 (ENCFF993IIG for peak and ENCFF699IRY for bigwig), H3K36me3 (ENCFF362DZS for peak and ENCFF703RVT for bigwig), H3K9me3 (ENCFF925BSH for peak and ENCFF293DGT for bigwig), H3K9ac (ENCFF668UBL for peak and ENCFF331XNQ for bigwig), CTCF (ENCFF533APC for peak and ENCFF954GDC for bigwig), POLR2A (ENCFF128LHX for peak and ENCFF291FGY for bigwig). For log2 fold change enrichment analysis, we computed the proportion of boundary imaged segments overlapped with ChIP-seq peaks, denoted as *p*_l_, and the proportion of all imaged segments overlapped with ChIP-seq peaks, denoted as *p*, and defined the log2 enrichment as *log*_2_*(p*_l_*/p)*. For Figure 3d analysis, we averaged the bigwig signal across all TAD boundaries for each of the 25 Kb interval within the 250 Kb window.

ChIP-seq data to validate IMR-90 TADs from Su et al^34^ is downloaded from ENCODE^49^ with the IDs: CTCF (ENCFF203SRF for peak and ENCFF101PSS for bigwig), POLR2A (ENCFF448ZOJ for peak and ENCFF444OMD for bigwig), SMC3 (ENCFF770ISZ for peak and ENCFF775AQV for bigwig), H3K4me3 (ENCFF093NQC for peak and ENCFF518GFI for bigwig), H3K36me3 (ENCFF449ADN for peak and ENCFF314QXF for bigwig), H3K9me3 (ENCFF098XMT for peak and ENCFF733CJA for bigwig), H3K9ac (ENCFF309ZMM for peak and ENCFF682HNO for bigwig).

### Single-cell TAD-like structures in mESCs by FISHnet

We ran FISHnet on autosomes from mESCs 25 Kb chromatin tracing data. Chromatin traces with more than 40% missingness are discarded, and the remaining traces are linearly imputed so that no 3D coordinates are missing. We then calculated the pairwise distance matrix for each chromatin and ran FISHnet with parameters `distance = [100, 150, 200, 250, 300,350, 400, 450, 500, 550, 600, 650]`, `plateau_size = 4`, `window_size = 2`, `size_exclusion = 3`, and `merge = 3`. For each imaged segment, we then computed how often it is marked as a boundary in individual traces.

### Hi-C A/B compartment enrichment

The bulk Hi-C A/B compartment assignment from Rao et al^10^ is downloaded from 4DN data portal with ID 4DNFIHM89EGL. For the log2 fold change enrichment, we computed the enrichment of different epigenetic markers in the A compartment. Specifically, we calculated the proportion of A compartments overlapped with ChIP-seq peaks, denoted as *p*_l_, and the proportion of all imaged segments overlapped with ChIP-seq peaks, denoted as *p*, and defined the log2 enrichment as *log*_2_*(p*_l_*/p)*.

### scRNA-seq data analysis

The scRNA-seq count matrix and cluster assignment from Loo et al^55^ is downloaded from the Github repository https://github.com/jeremymsimon/MouseCortex with file names E14_combined_matrix.txt.gz and E14_combined_matrix_ClusterAnnotations.txt. The count matrix is filtered by removing cells that contain more than 10% mitochondrially derived transcripts, removing cells with fewer than 500 genes detected, and removing genes present in fewer than 10 cells or having fewer than 60 total transcripts. After the filtered count matrix is normalized and log-transformed, the combat function from ScanPy^70^ is used to correct for batch effects. The cell clusters with high *Neurod6* expression are treated as excitatory neurons. For relative expression of *Synj1*, we downloaded a list of housekeeping genes from https://github.com/brianpenghe/Matlab-genomics, with file name He_2020_ENCODE3_RNA/GeneLists/Bulk Cluster Ubiquitous.txt. For each single cell, we subtract the log-transformed expression of *Synj1* by the average log-transformed expression of housekeeping genes in that cell and calculated the average log fold change of expression across all cells. The scRNA-seq data from Guo et al is already normalized, so we only log-transformed it and computed the log fold change of expression.

## Data availability

Chromatin tracing datasets from Takei et al^23^, Huang et al^52^, and Takei et al^24^ are downloaded from the 4DN data portal: 25 Kb DNA seqFISH+ mouse embryonic stem cells data (4DNFIHF3JCBY for the first biological replicate and 4DNFIQXONUUH for the second biological replicate), 5 Kb multiplexed DNA FISH data (4DNFI2RCYFJU), and 25 Kb DNA seqFISH+ mouse brain cells data (4DNFIW4S8M6J for the first biological replicate, 4DNFI4LI6NNV for the second biological replicate, and 4DNFIDUJQDNO for the third biological replicate). If the data contains multiple 3D coordinates for the same segment, only the first is kept. Chromatin tracing data from Su et al^34^ are downloaded from https://zenodo.org/records/3928890.

## Code availability

The ArcFISH software is available at https://github.com/hyuyu104/ArcFISH. Documentation is available at https://hyuyu104.github.io/ArcFISH/. Source code to reproduce analysis is available at https://github.com/hyuyu104/ArcFISH/tree/main/figures.

## Acknowledgements

M.H. was partially supported by the NIH grants R35HG011922 and UM1HG011585. Y.L. was partially supported by R01AG079291 and P50HD103573.

## Author contributions

H.Y., M.H., and Y.L. conceived the project. H.Y. and L.Z. analyzed the data. H.Y., M.H., and Y.L. wrote the manuscript with input from all other authors.

