## Supplementary File for "Accurate and robust 3D genome feature discovery from multiplexed DNA FISH"

#### Contents

**1 Supplementary Figures** **2**

**Supplementary Figure 1** **2**
**Supplementary Figure 2** **3**
**Supplementary Figure 3** **4**
**Supplementary Figure 4** **5**
**Supplementary Figure 5** **6**
**Supplementary Figure 6** **7**
**Supplementary Figure 7** **8**
**Supplementary Figure 8** **9**
**Supplementary Figure 9** **10**
**Supplementary Figure 10** **11**
**Supplementary Figure 11** **12**
**Supplementary Figure 12** **13**
**Supplementary Figure 13** **14**
**Supplementary Figure 14** **15**
**Supplementary Figure 15** **16**
**Supplementary Figure 16** **17**
**Supplementary Figure 17** **18**
**Supplementary Figure 18** **19**
**Supplementary Figure 19** **20**
**Supplementary Figure 20** **21**

**2 Supplementary methods** **22**

### 1 Supplementary Figures

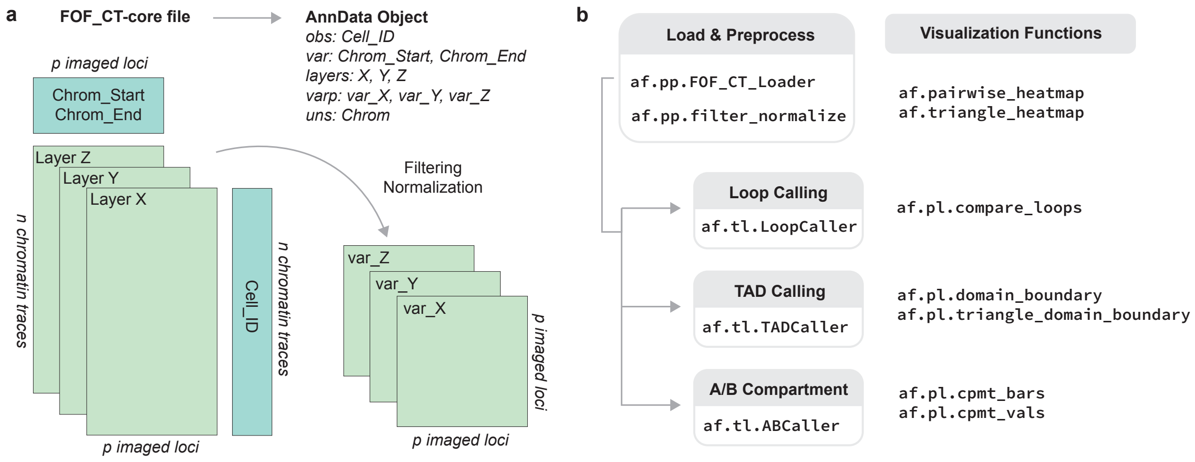

**Supplementary Figure 1**. **Illustration of ArcFISH software.** **a**, FOF_CT-core files are stored as AnnData objects in ArcFISH. Each AnnData object stores the information of one chromosome, indicated by the Chrom field in uns. Layers X, Y, and Z store the 3D coordinates of chromatin traces of this chromosome. Only the axis variance matrices, var_X, var_Y, and var_Z, are stored instead of the entire pairwise difference tensor. **b**, Main utilities in ArcFISH python software.

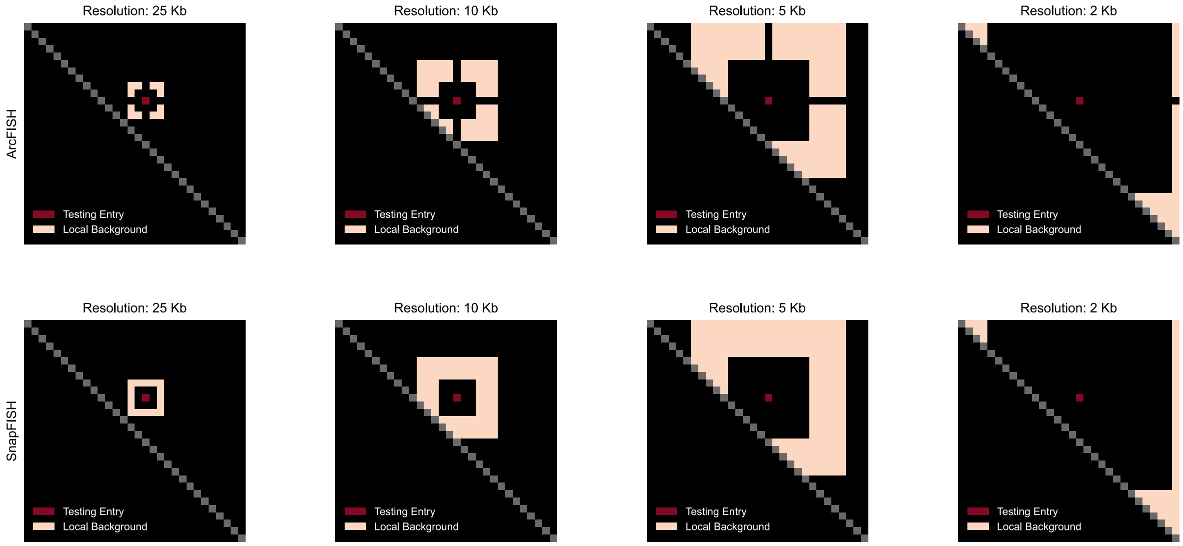

**Supplementary Figure 2**. **Chromatin loop local background in SnapFISH and ArcFISH.** The red pixel represents the entry currently testing, and the white entries are the local background. We only consider the upper triangle. The plots show example regions with 30 loci at resolutions 25Kb, 10Kb, 5Kb, and 2Kb. The pixel currently testing is the (10, 16) locus pair.

**a**

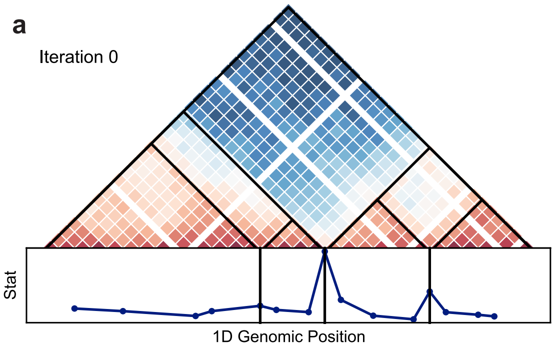

| c1 | s1 | e1 | c2 | s2 | e2 | level |
|---|---|---|---|---|---|---|
| chr2 | 109000000 | 109025000 | chr2 | 109675000 | 109700000 | 0 |
| chr2 | 109675000 | 109700000 | chr2 | 109875000 | 109900000 | 0 |
| chr2 | 109875000 | 109900000 | chr2 | 110200000 | 110225000 | 0 |
| chr2 | 110200000 | 110225000 | chr2 | 110525000 | 110550000 | 0 |

**b**

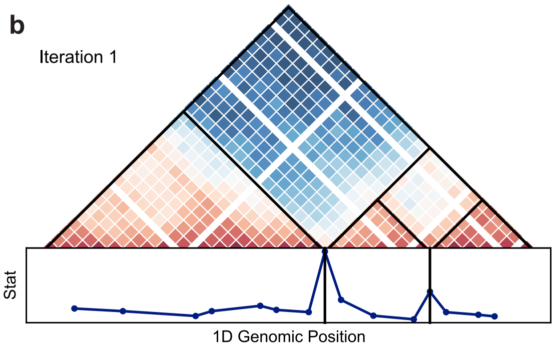

| c1 | s1 | e1 | c2 | s2 | e2 | level |
|---|---|---|---|---|---|---|
| chr2 | 109000000 | 109025000 | chr2 | 109675000 | 109700000 | 0 |
| chr2 | 109675000 | 109700000 | chr2 | 109875000 | 109900000 | 0 |
| chr2 | 109875000 | 109900000 | chr2 | 110200000 | 110225000 | 0 |
| chr2 | 110200000 | 110225000 | chr2 | 110525000 | 110550000 | 0 |
| chr2 | 109000000 | 109025000 | chr2 | 109875000 | 109900000 | 1 |

**c**

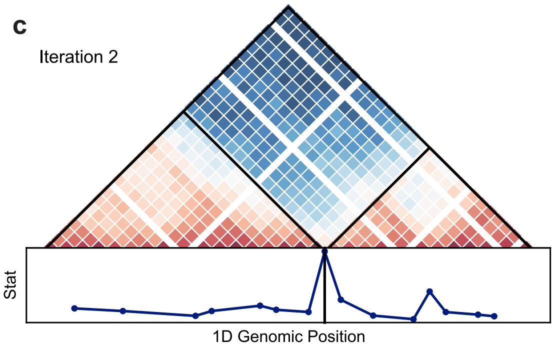

| c1 | s1 | e1 | c2 | s2 | e2 | level |
|---|---|---|---|---|---|---|
| chr2 | 109000000 | 109025000 | chr2 | 109675000 | 109700000 | 0 |
| chr2 | 109675000 | 109700000 | chr2 | 109875000 | 109900000 | 0 |
| chr2 | 109875000 | 109900000 | chr2 | 110200000 | 110225000 | 0 |
| chr2 | 110200000 | 110225000 | chr2 | 110525000 | 110550000 | 0 |
| chr2 | 109000000 | 109025000 | chr2 | 109875000 | 109900000 | 1 |
| chr2 | 109875000 | 109900000 | chr2 | 110525000 | 110550000 | 2 |

**Supplementary Figure 3**. **Illustration of hierarchical TAD calling.** **a**, The raw boundary calling result. Vertical lines show all three TAD boundaries identified by ArcFISH. The three boundaries partition the imaging region to four TADs, which are appended to the output TAD list with level=0. **b**, TAD list after one iteration. In the first iteration, the boundary with the smallest statistic is removed, creating a new TAD. This TAD is then added to the TAD list with level=1. **c**, TAD list after two iterations. By the same procedure, a new TAD is added with level=2.

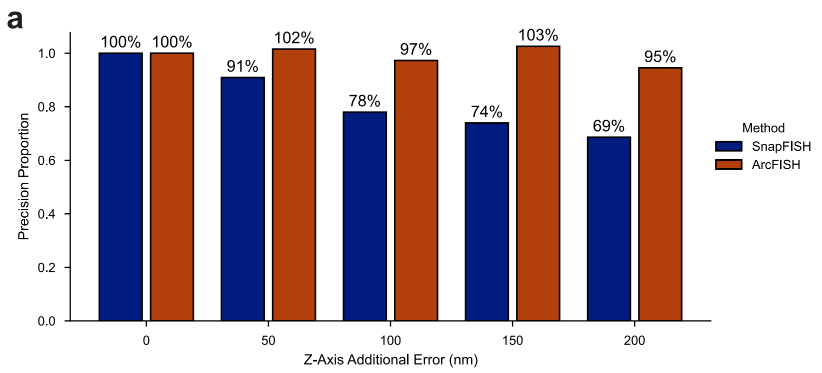

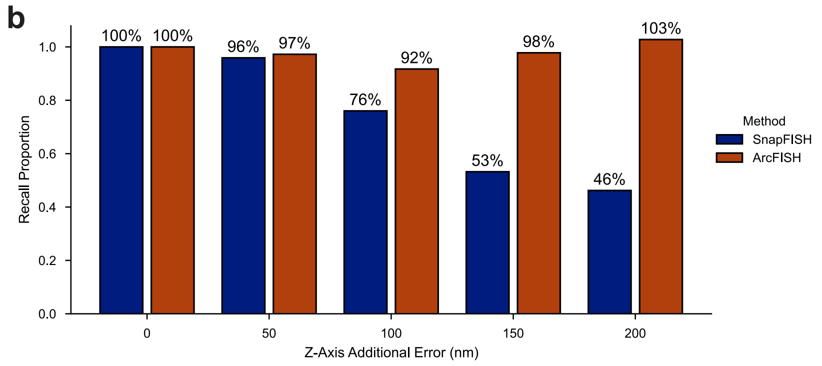

**Supplementary Figure 4**. **Precision and recall across different noise levels as a fraction of uniform noise data.** **a**, Average precision as a fraction of uniform noise data. **b**, Average recall as a fraction of uniform noise data. The precision and recall are averaged from n=10 replicates. The bar at each noise level denotes the precision or recall at this noise level as a fraction of the ones from 0nm Z-axis noise data.

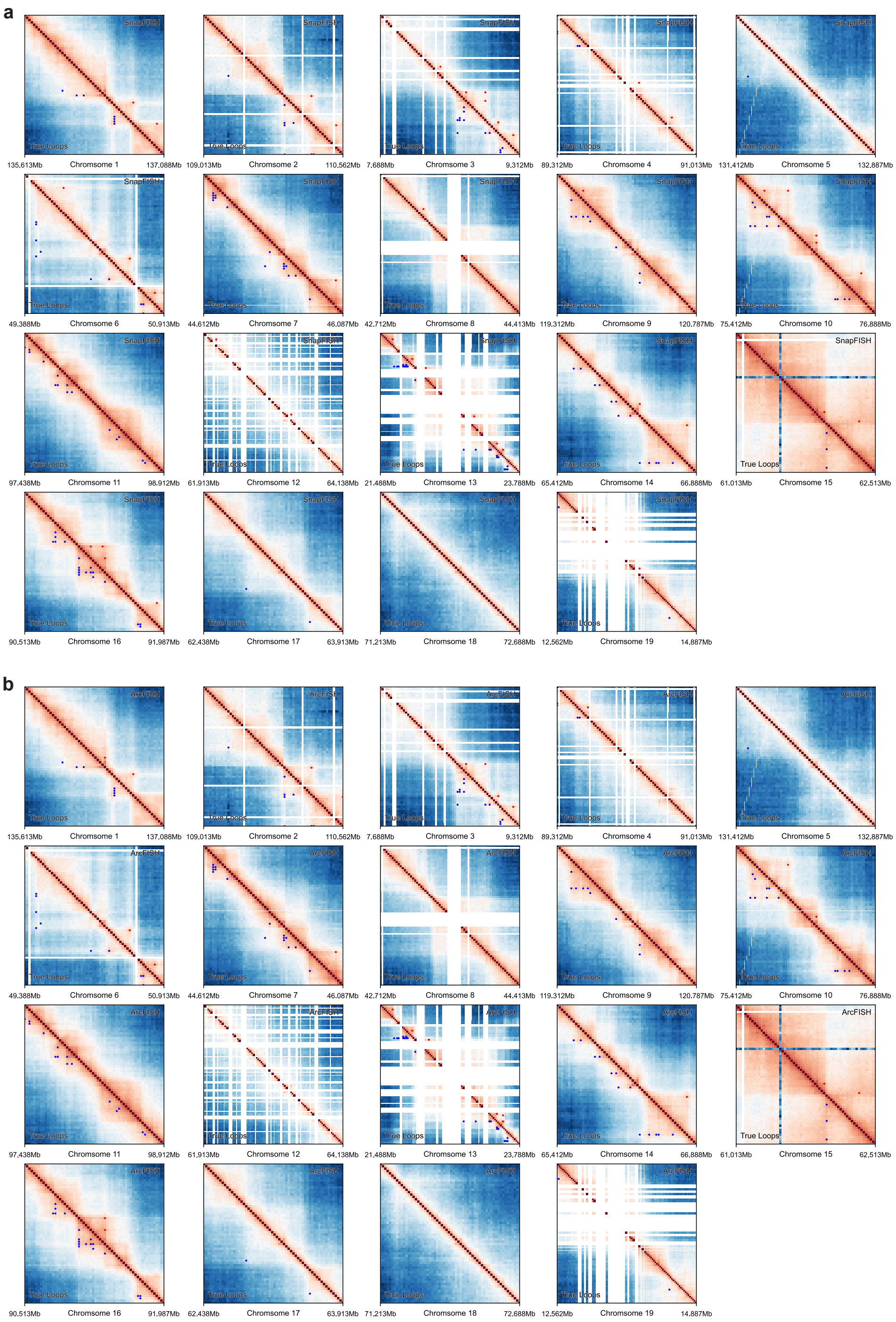

**Supplementary Figure 5**. **Loop calling results of DNA seqFISH+ mESC 25Kb data.** **a**, Loop calling results from SnapFISH. **b**, Loop calling results from ArcFISH. Heatmaps show the median pairwise distance matrix in nm. Blank areas are positions not covered by seqFISH+ as the loci are not adjacent to each other in some chromosomes. Blue dots are the positions of true loops, i.e., loops present in at least two of the six reference lists. Red dots are the positions of SnapFISH (in **a**) or ArcFISH (in **b**) loops.

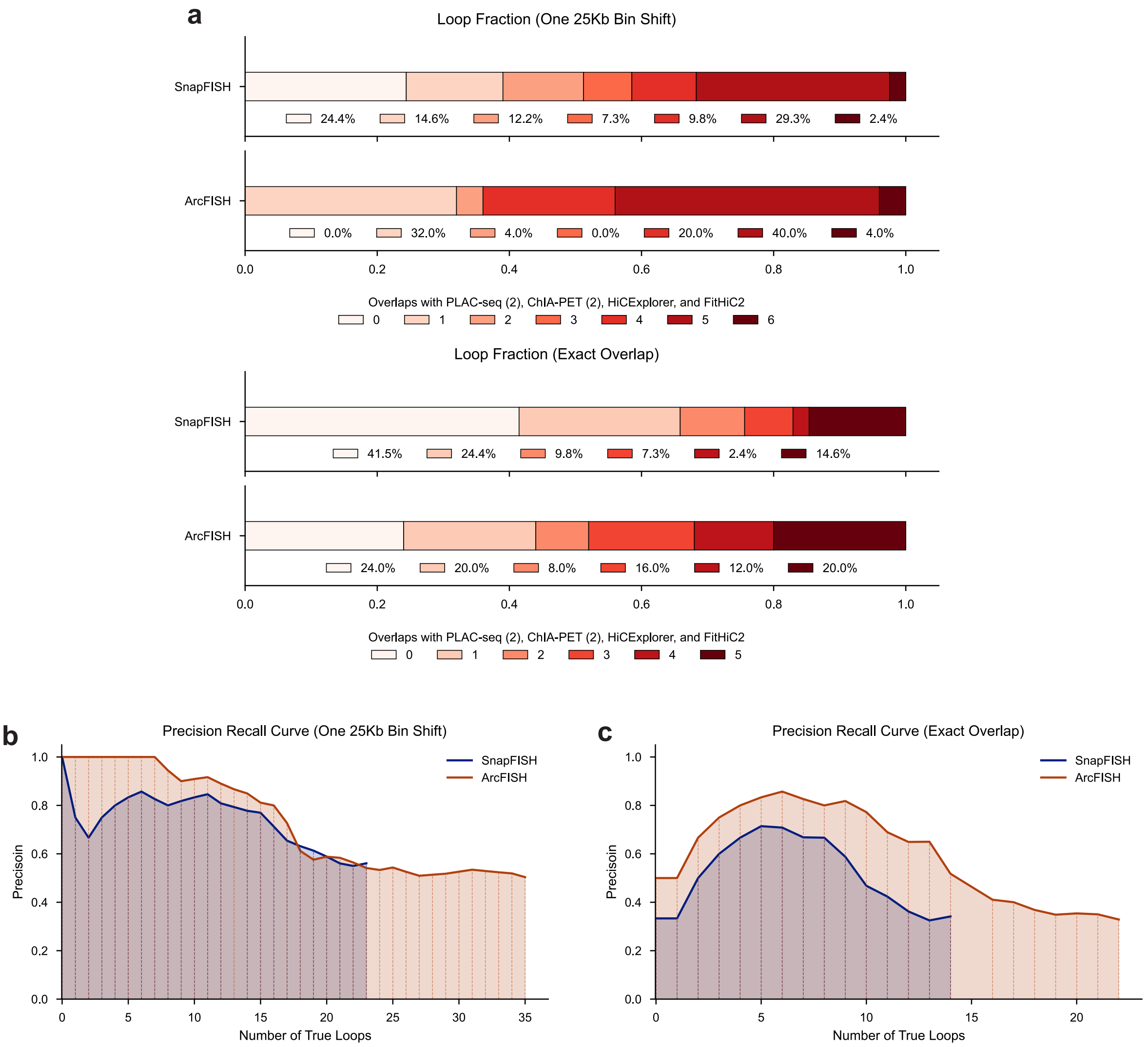

**Supplementary Figure 6**. **ArcFISH has higher overlap with the reference loop lists regardless of p-value cutoffs. a**, Fractions of SnapFISH and ArcFISH loops overlapped with reference loop lists. The six reference loop lists are MAPS CTCF PLAC-seq, MAPS H3K4me3 PLAC-seq, CTCF ChIA-PET, POLR2A ChIA-PET, HiCExplorer, and FitHiC2. In the top panel, two loops are considered overlapped if both loop anchors are within 25Kb of each other. In the bottom panel, two loops are considered overlapped if they have the exact same loop anchors. **b**, Precision recall curve allowing 25Kb shift. The curve is obtained by varying the final p-value cutoff of both SnapFISH and ArcFISH. At each cutoff, loops identified are labeled as true positives if they have both anchors within 25Kb of a loop from the true loop list. **c**, Precision recall curve requiring exact overlap. It is similar to **b** but counts true positives based on exact overlap.

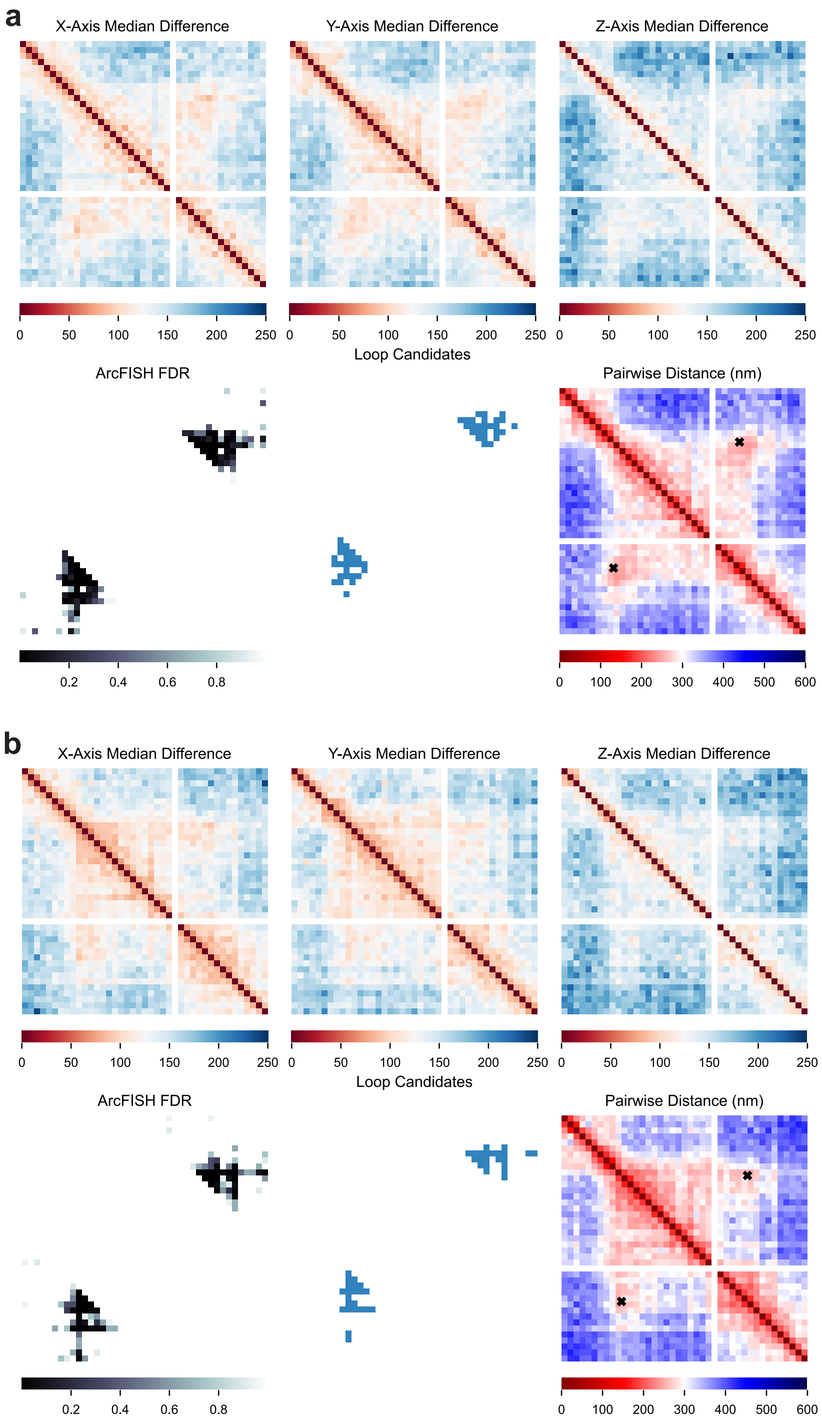

**Supplementary Figure 7**. **Loop identification from multiplexed DNA FISH 5Kb mESC data.** **a**, ArcFISH identifies one chromatin loop from the 129 allele. The top row shows the median pairwise difference from each axis. The second row shows the FDR, loop candidates, and chromatin loop outputted by ArcFISH overlaid on the median pairwise distance matrix. **b**, ArcFISH identifies one chromatin loop from the CAST allele. The panels are organized in the same order as in **a**.

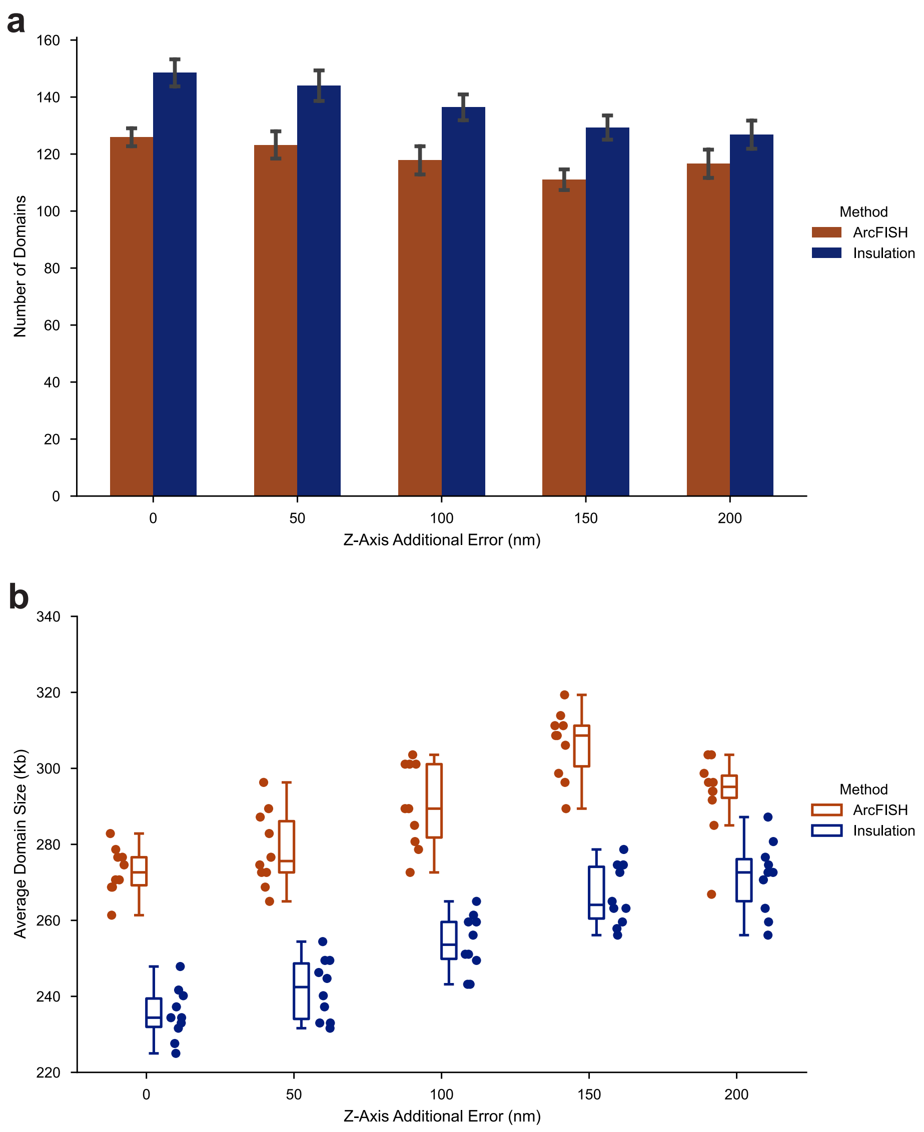

**Supplementary Figure 8**. **Number and size of domains called by ArcFISH and Insulation Score across different noise levels. a**, Number of domains across different noise levels. Error bar represents the standard deviation computed from n=10 replicates. **b**, Domain sizes across different noise levels. n=10 replicates are generated for each noise level.

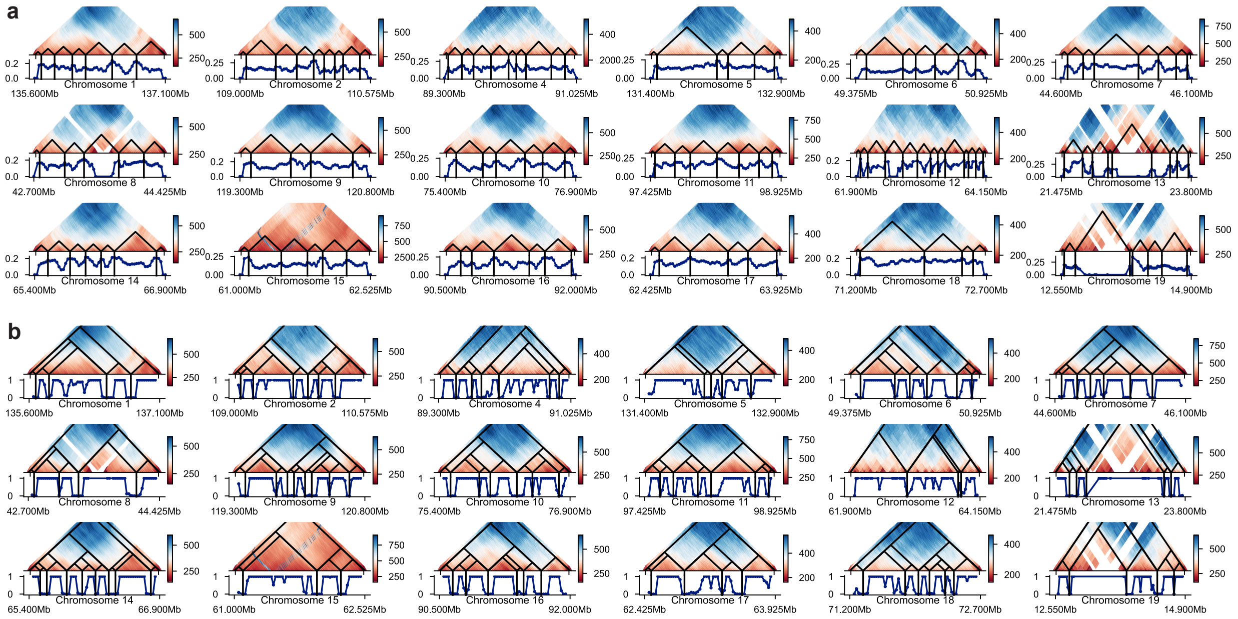

**Supplementary Figure 9**. **Domain calling results of DNA seqFISH+ mESC 25Kb data. a**, Domain calling results from Insulation Score. Chromosome 3 is omitted as it is already shown in Figure 3c. For each chromosome, the heatmap is the median pairwise distance matrix in nm, and the line plot shows the insulation score. **b**, Domain calling results from ArcFISH. The hierarchical structure of TADs and subTADs is also displayed in the heatmaps. The line plot shows the FDR from ArcFISH.

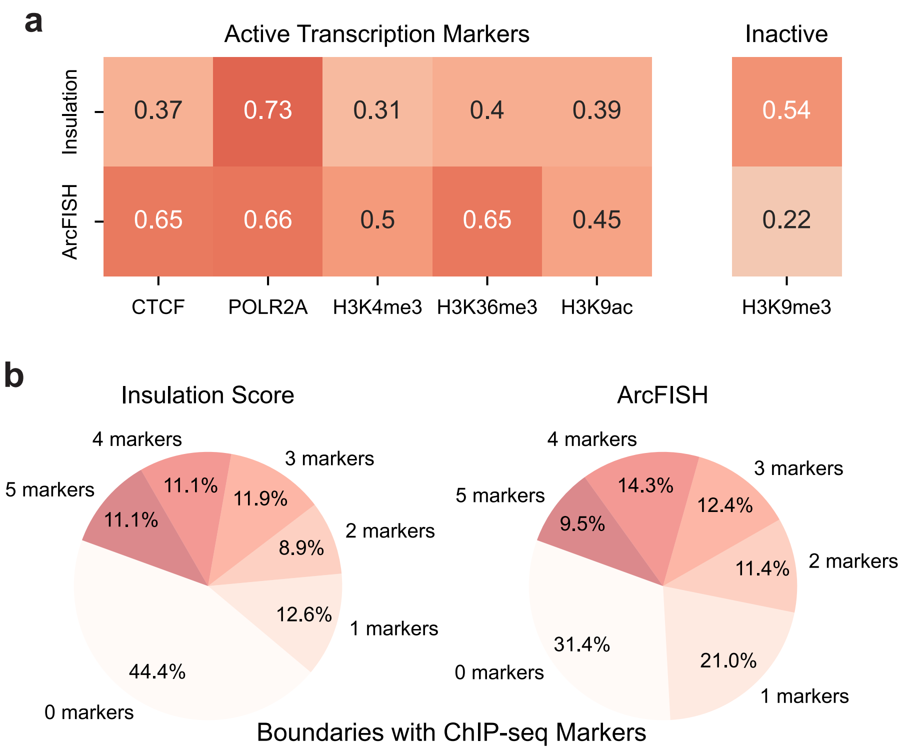

**Supplementary Figure 10**. **ArcFISH boundaries show better correspondence with epigenetic markers. a**, Enrichment of ChIP-seq peaks in boundaries. The first five columns are markers for active transcription, thus expected to be enriched at boundaries. The last column is a marker for inactive transcription and thus should be depleted at boundaries. Values are the log2 fold changes between the boundary and the entire imaging region. **b**, Fraction of boundaries overlapping with active transcription markers. The five markers are CTCF, POLR2A, H3K4me3, H3K36me3, and H3K9ac.

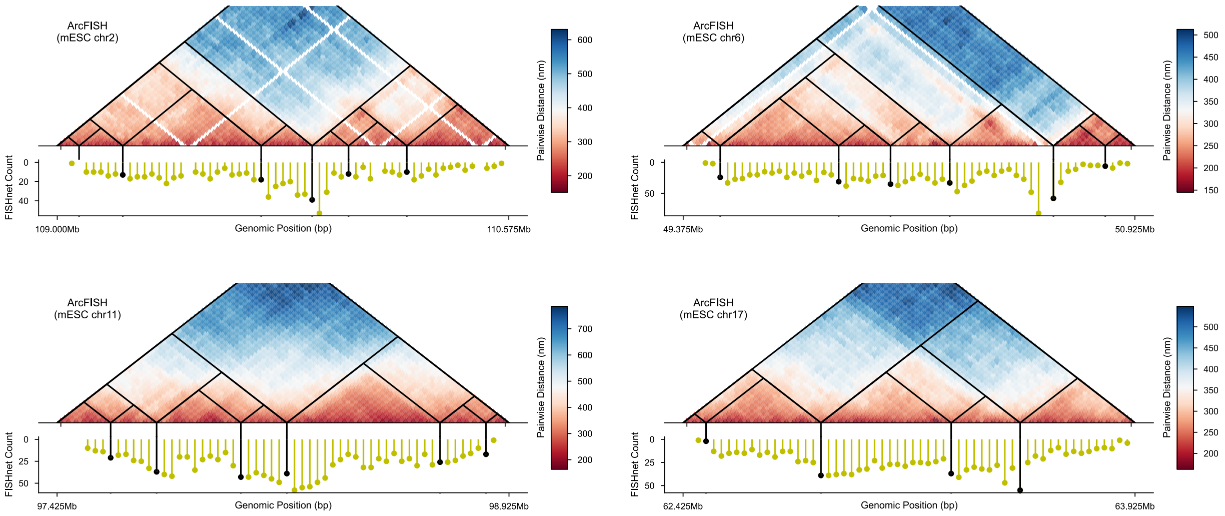

**Supplementary Figure 11**. **ArcFISH boundaries appear frequently in single cells.** Example heatmaps from chromosomes 2, 6, 11, and 17 with ArcFISH TADs overlaid. The dot plot below displays how many times a locus is identified as a boundary by FISHnet in single traces. The black lines correspond to the count of ArcFISH boundaries.

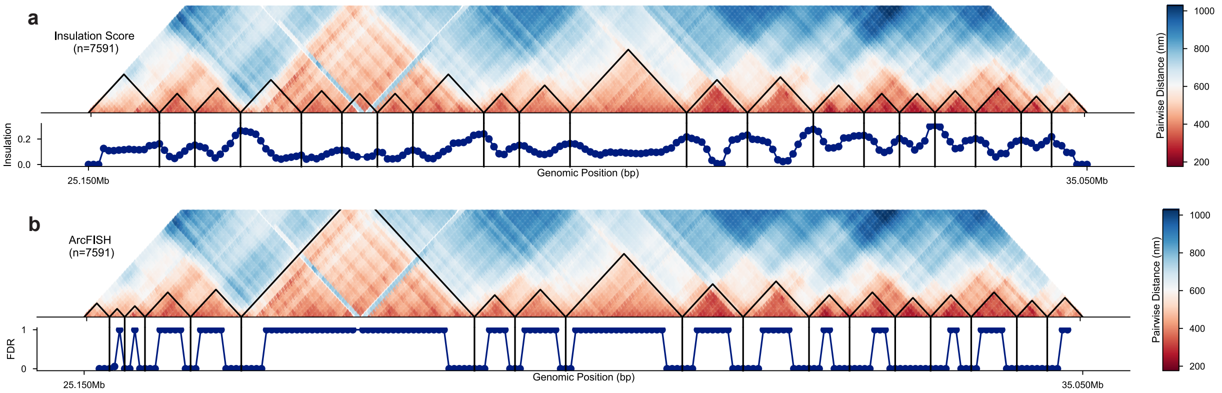

**Supplementary Figure 12**. **TAD identified from chromosome 21. a**, Insulation Score identified TADs in an example region. **b**, ArcFISH identified TADs in an example region.

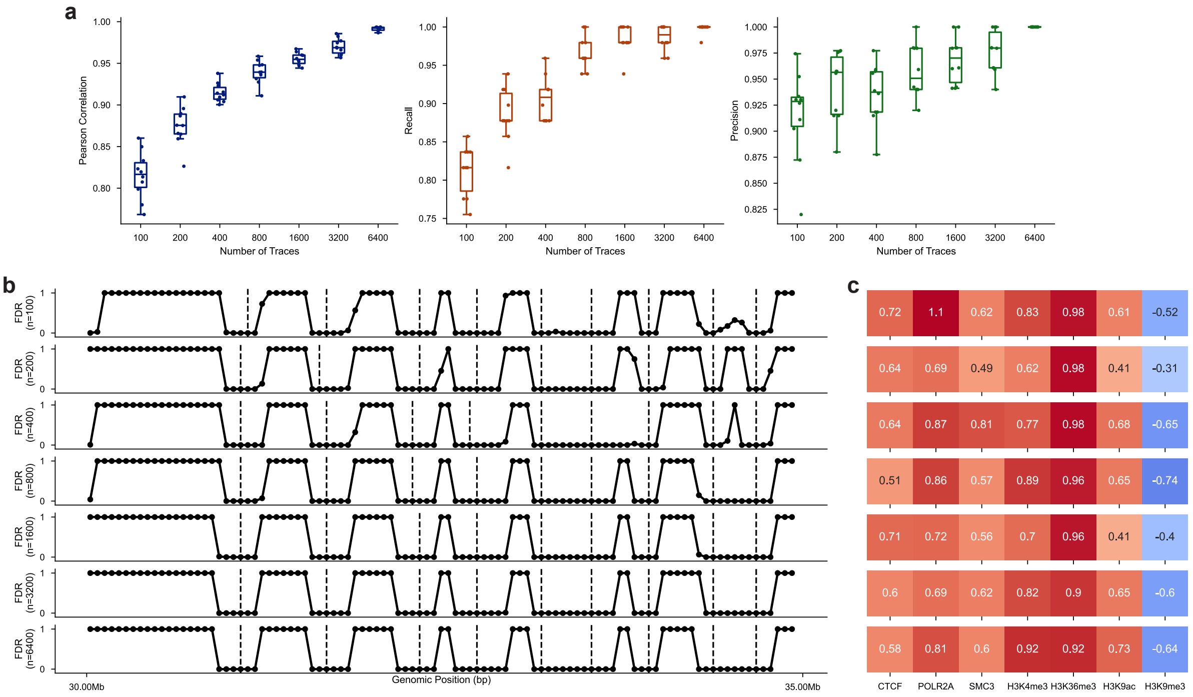

**Supplementary Figure 13. Domain calling performance with different number of traces. a**, FDR correlation, precision, and recall with different number of traces. FDR correlation is computed from the full data and the sampled data. Precision and recall are calculated by treating the boundaries from the full data as the ground truth. n=10 replicates are generated for each number of traces. **b**, Example FDR profile for different number of traces. **c**, Enrichment of different epigenetic markers at domain boundaries across different number of traces.

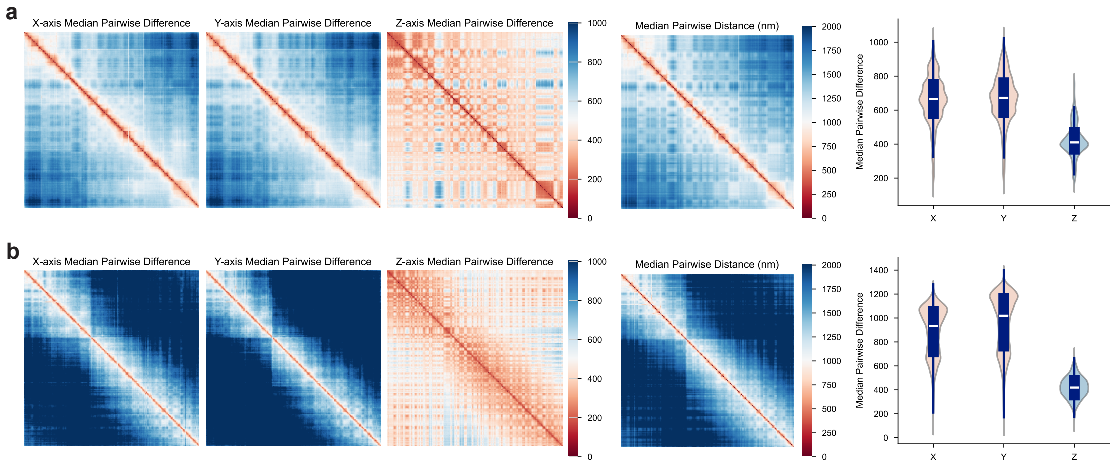

**Supplementary Figure 14**. **Chromosome 2 p-arm and entire chromosome heatmaps from Su et al.** **a**, Z-axis shows different noise level in p-arm. The heatmaps are median pairwise difference matrices for each axis and median pairwise distance matrix. The violin plot shows the distribution of median pairwise difference, which should be similar for X, Y, and Z axes if their noise levels are the same. **b**, Z-axis shows different noise level in entire chromosome 2. Plots are organized in the same order as in **a**. Still, from the median pairwise difference and the violin plot, Z-axis is different from X and Y axes.

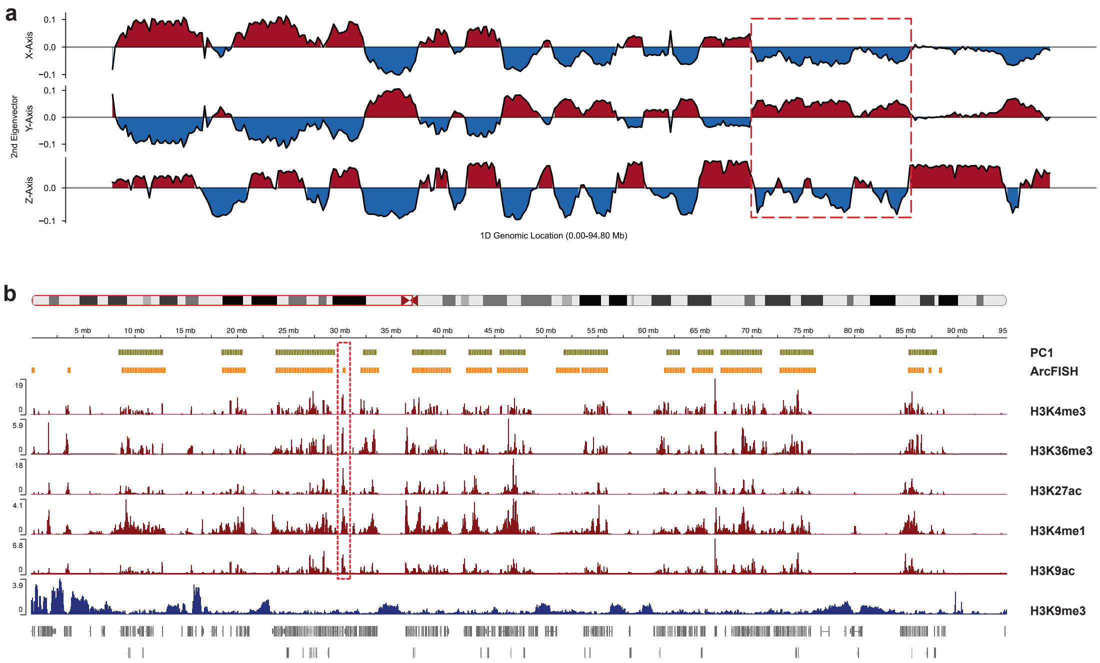

**Supplementary Figure 15**. **ArcFISH compartment assignments of chromosome 2 p-arm. a**, Chromosome 2 p-arm second eigenvectors from each axis. The red dashed box highlights a region where the X and Y axes fail to distinguish A/B compartments while Z-axis has clear alternating signals. **b**, PC1 and ArcFISH A compartments correlate with transcription markers. The top two tracks show the A compartments from PC1 and ArcFISH. The red tracks are active transcription markers, and the blue track is an inactive transcription marker. The red box shows a region where ArcFISH A compartment coincides with peaks from active transcription marker but PC1 fails.

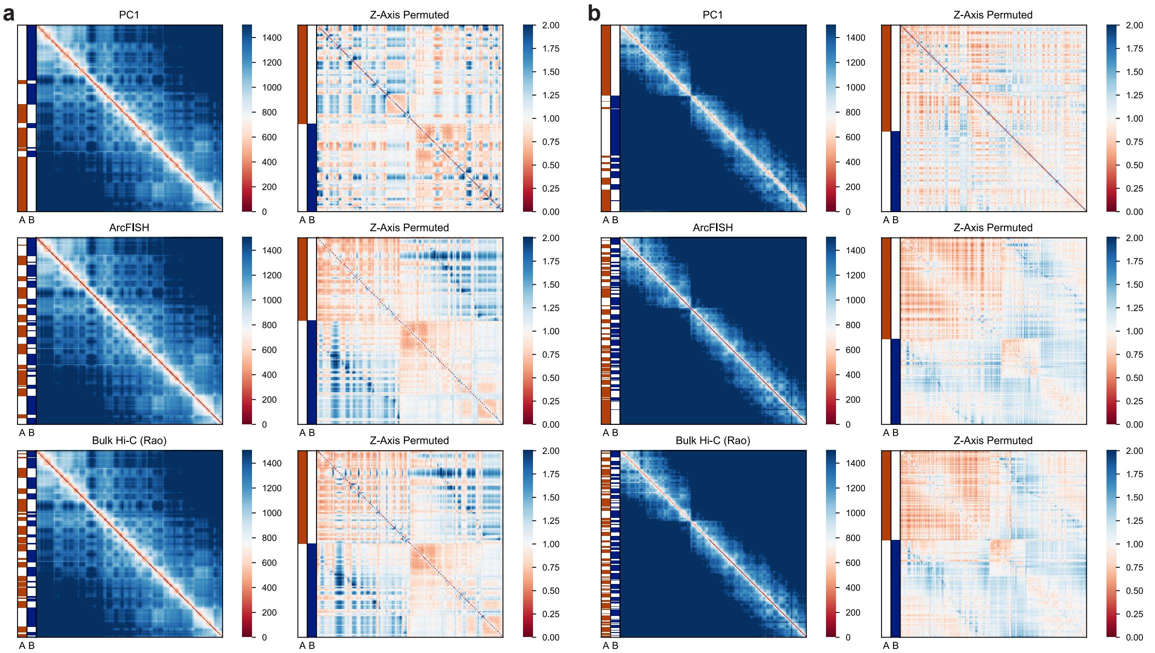

**Supplementary Figure 16**. **Compartment assignments from 30 traces p-arm and 3029 traces entire chromosome 2**. **a**, A/B compartment assignments of 30 traces p-arm. The first column shows the compartment assignment juxtaposed to the median pairwise distance matrix from the original data (4848 traces). The second column shows the Z-axis pairwise difference permuted according to the compartment assignments. As A and B compartments tend to interact more frequently within than between, the permuted heatmap is expected to show a two-block pattern. **b**, A/B compartment assignments of entire chromosome 2. Plots are organized in the same way as in **a**.

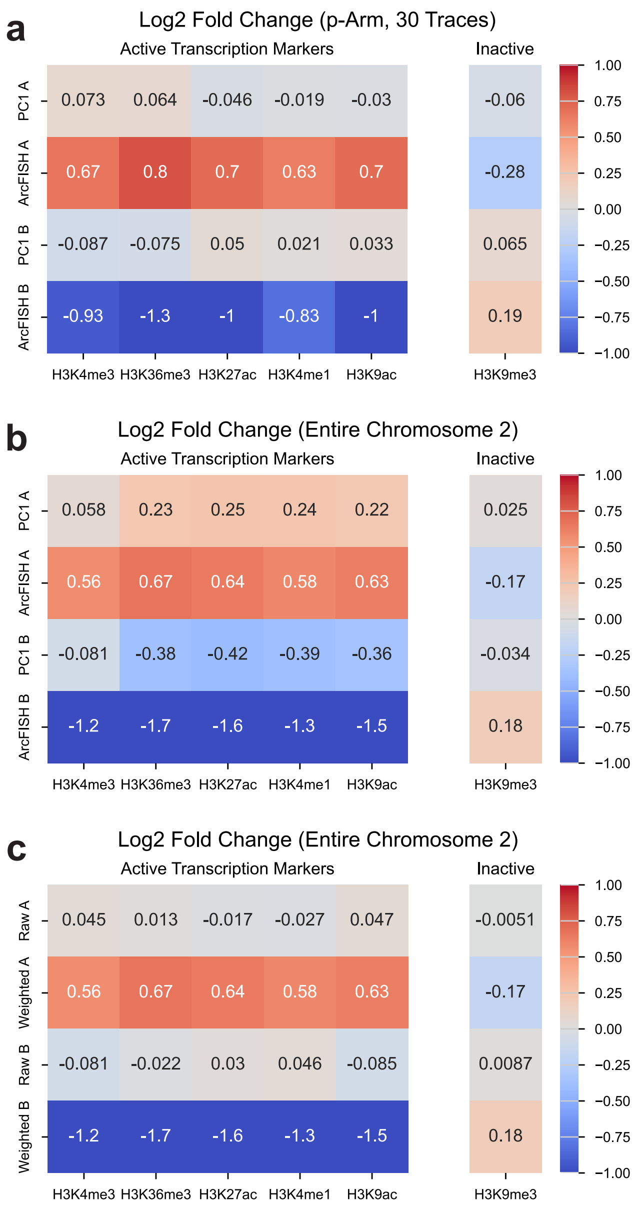

**Supplementary Figure 17**. **Enrichment of epigenetic markers in A/B compartments under different scenarios.** **a**, Log2 fold change of 30 traces chromosome 2 p-arm. **b**, Log2 fold change of entire chromosome from PC1 and ArcFISH. **c**, Log2 fold change of entire chromosome from unweighted second eigenvectors and weighted second eigenvector (ArcFISH).

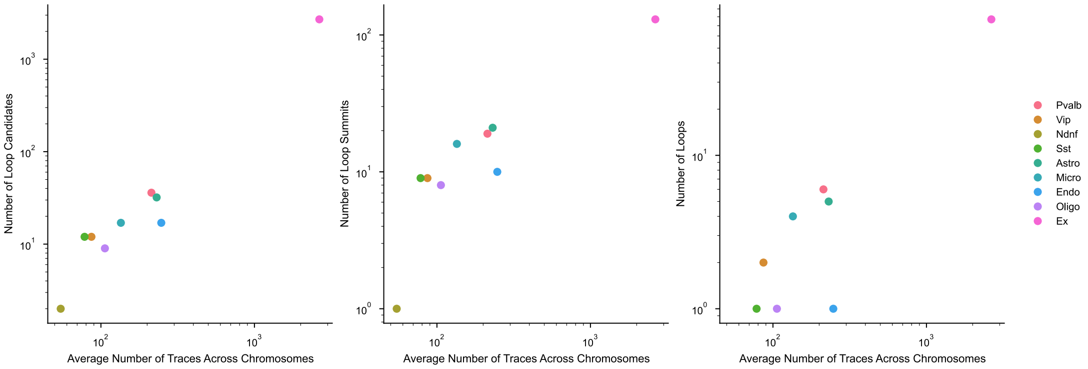

**Supplementary Figure 18**. **Number of loops increases with the number of traces.** The number of loop candidates, loop summits, and final loops versus the number of chromatin traces.

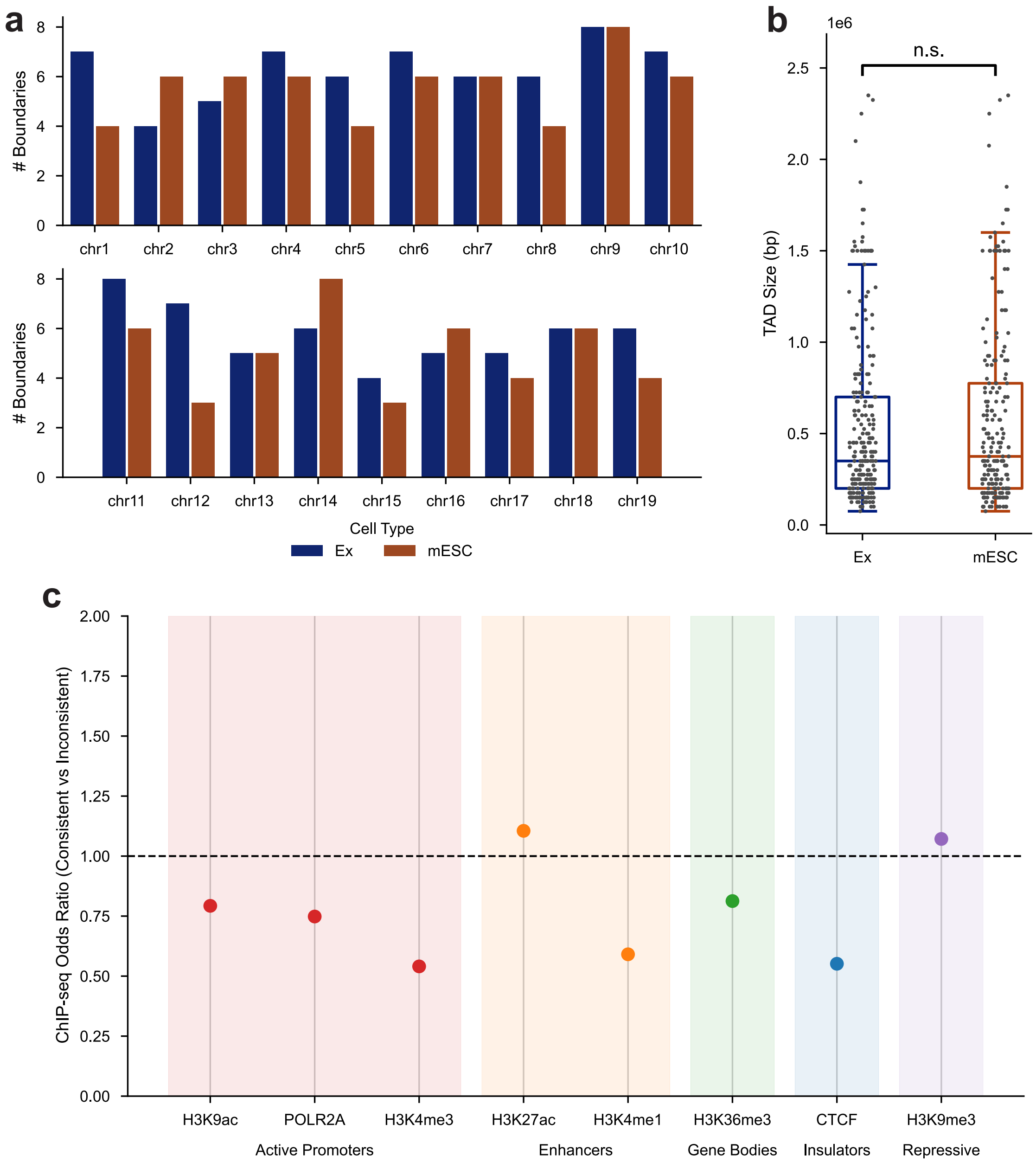

**Supplementary Figure 19**. **Enrichment of epigenetic markers in A/B compartments under different scenarios. a**, Number of TAD boundaries in excitatory neurons and mESCs by chromosome number. **b**, Excitatory neurons and mouse embryonic stem cells have similar TAD size. **c**, Histone mark odds ratio between shared and cell-type-specific TAD boundaries. Histone marks are colored based on their biological interpretation: H3K9ac, POLR2A, and H3K4me3 label active promotors; H3K27ac and H3K4me1 label active enhancers; H3K36me3 labels gene bodies; CTCF labels insulators; and H3K9me3 is a repressive transcription marker.

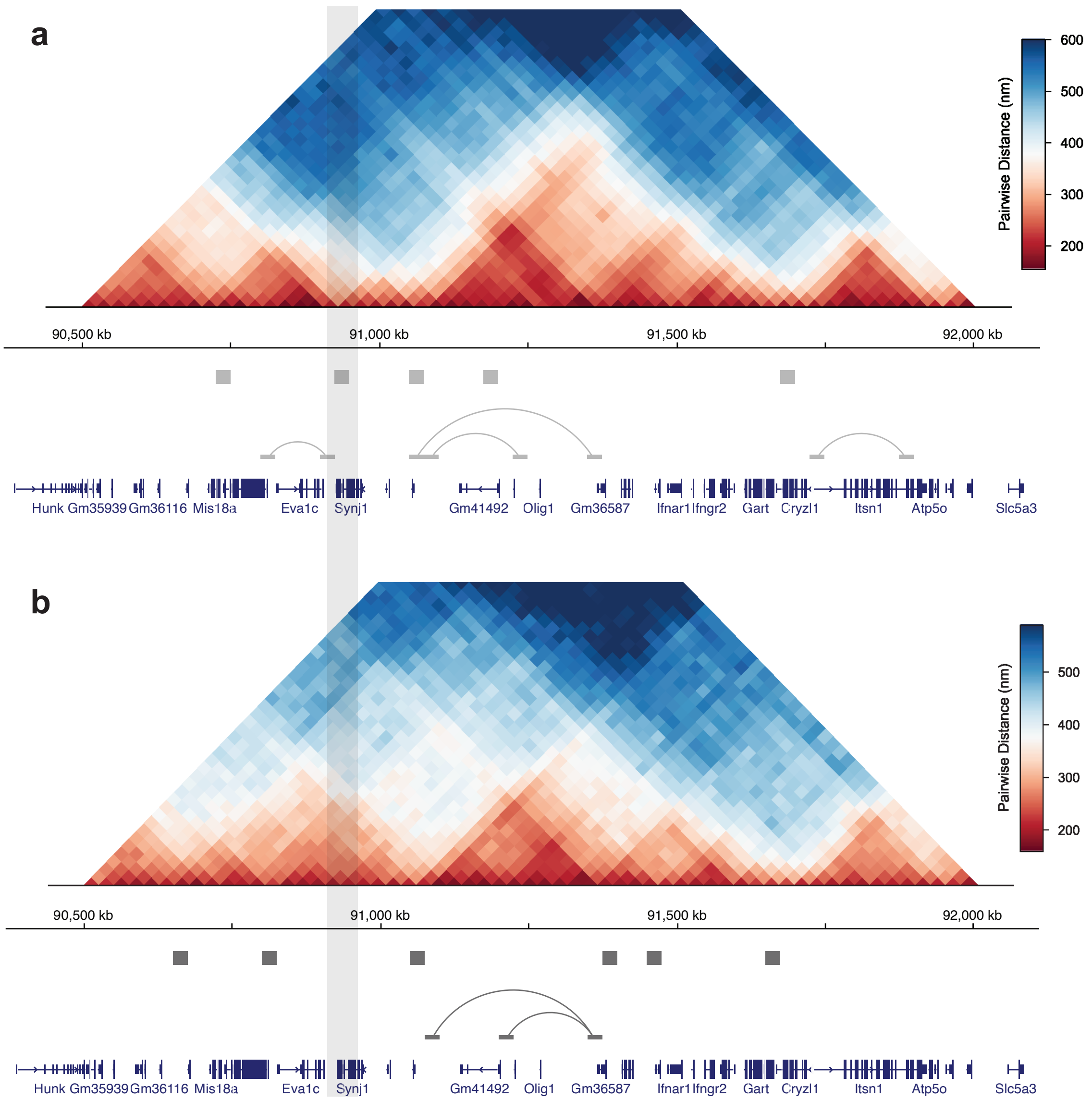

**Supplementary Figure 20. Pairwise distance heatmaps of chromosome 16. a**, Excitatory neurons median pairwise distance. **b**, mESCs median pairwise distance. Gray box highlights the position of *Synj1* gene.

### 2 Supplementary methods

#### 2.1 Identifiability of measurement errors

We first explain why the measurement errors $\sigma_x^2$, $\sigma_y^2$, and $\sigma_z^2$ are not identifiable under the original model assumptions. The statistical model for the chromatin trace, as defined in Methods, is

$$\begin{pmatrix}\mathbf{x}\\ \mathbf{y}\\ \mathbf{z}\end{pmatrix} \sim \mathrm{N}\left(\mathbf{0}, \begin{bmatrix}\mathbf{K}+\sigma_x^2\mathbf{I}_p & & \\ & \mathbf{K}+\sigma_y^2\mathbf{I}_p & \\ & & \mathbf{K}+\sigma_z^2\mathbf{I}_p\end{bmatrix}\right),$$

i.e., each axis has a Gaussian distribution with the same chromatin conformation but different measurement error, and the axes are independent of each other. To show that the measurement errors are not identifiable, we need to find two distinct sets of parameters $(\sigma_x^2, \sigma_y^2, \sigma_z^2, \mathbf{K})$ and $(\tau_x^2, \tau_y^2, \tau_z^2, \mathbf{S})$ that give the same distribution of $(\mathbf{x}^\top, \mathbf{y}^\top, \mathbf{z}^\top)$. Moreover, these two parameter sets should lie in the parameter space: $\mathbf{K}, \mathbf{S} \succeq 0$ (i.e., positive semidefinite) and all measurement errors are nonnegative.

Indeed, for any nonzero $(\sigma_x^2, \sigma_y^2, \sigma_z^2)$ and any positive semidefinite $\mathbf{K}$, define the positive constant $c = \frac{1}{2}\min\{\sigma_x^2, \sigma_y^2, \sigma_z^2\}$. Let $\mathbf{S} = \mathbf{K} + c\mathbf{I}_p$ and

$$\begin{cases}\tau_x^2 = \sigma_x^2 - c\\ \tau_y^2 = \sigma_y^2 - c\\ \tau_z^2 = \sigma_z^2 - c\end{cases}.$$

Then $\tau_x^2, \tau_y^2, \tau_z^2 > 0$ and $\mathbf{S} \succeq 0$ so that they lie in the parameter space, and the two sets of parameters are different: $(\sigma_x^2, \sigma_y^2, \sigma_z^2, \mathbf{K}) \neq (\tau_x^2, \tau_y^2, \tau_z^2, \mathbf{S})$. Moreover, they define the same Gaussian distribution as the means are both $\mathbf{0}$ and the covariance matrices are the same:

$$\mathrm{Blk\,diag}(\mathbf{S} + \tau^2\mathbf{I}_p) = \mathrm{Blk\,diag}(\mathbf{K} + c\mathbf{I}_p + \sigma^2\mathbf{I}_p - c\mathbf{I}_p) = \mathrm{Blk\,diag}(\mathbf{K} + \sigma^2\mathbf{I}_p).$$

Therefore, the measurement errors are not identifiable.

We next show how the parameters become identifiable after assuming the true squared pairwise distance matrix, $\tilde{\mathbf{D}}$, has rank less than $p/2 - 2$. The $ij$th entry of $\tilde{\mathbf{D}}$ is defined as

$$\tilde{d}_{ij} = \mathrm{E}\left[(\tilde{x}_i - \tilde{x}_j)^2 + (\tilde{y}_i - \tilde{y}_j)^2 + (\tilde{z}_i - \tilde{z}_j)^2\right],$$

which is the expected squared Euclidean distance between the $i$th locus and the $j$th locus when there are no measurement errors. For any fixed parameters $(\sigma_x^2, \sigma_y^2, \sigma_z^2, \mathbf{K})$,

$$\tilde{d}_{ij} = (K_{ii} + K_{jj} - 2K_{ij})\mathrm{E}\left[\chi_3^2\right] = 3(K_{ii} + K_{jj} - 2K_{ij})$$

since $\tilde{x}_i - \tilde{x}_j$, $\tilde{y}_i - \tilde{y}_j$, and $\tilde{z}_i - \tilde{z}_j$ are independent and identically distributed as normal with mean zero and variance $\mathrm{Var}(\tilde{x}_i - \tilde{x}_j) = \mathrm{Var}(x_i) + \mathrm{Var}(x_j) - 2\mathrm{Cov}(x_i, x_j) = K_{ii} + K_{jj} - 2K_{ij}$. In matrix form, this can be rewritten as $\tilde{\mathbf{D}} = 3(\mathbf{1}\mathbf{k}^\top + \mathbf{k}\mathbf{1}^\top - 2\mathbf{K})$, where $\mathbf{k} \in \mathbb{R}^p$ is the vector of diagonal elements of $\mathbf{K}$. Thus, $\mathbf{K} = \mathbf{1}\mathbf{k}^\top/2 + \mathbf{k}\mathbf{1}^\top/2 - \tilde{\mathbf{D}}/6$, and by the fact that $\mathrm{rank}(\mathbf{A} + \mathbf{B}) \leq \mathrm{rank}(\mathbf{A}) + \mathrm{rank}(\mathbf{B})$,

$$\mathrm{rank}(\mathbf{K}) \leq \mathrm{rank}(\mathbf{1}\mathbf{k}^\top/2) + \mathrm{rank}(\mathbf{k}\mathbf{1}^\top/2) + \mathrm{rank}(\tilde{\mathbf{D}}/6) < 2 + p/2 - 2 = p/2.$$

Therefore, imposing a low-rank assumption on $\tilde{\mathbf{D}}$ makes $\mathbf{K}$ not of full rank as well.

Now suppose $(\sigma_x^2, \sigma_y^2, \sigma_z^2, \mathbf{K})$ and $(\tau_x^2, \tau_y^2, \tau_z^2, \mathbf{S})$ are two sets of parameters leading to the same distribution. As their corresponding covariance matrices are the same,

$$\mathbf{K} + \sigma_x^2 \mathbf{I}_p = \mathbf{S} + \tau_x^2 \mathbf{I}_p \quad \mathbf{K} + \sigma_y^2 \mathbf{I}_p = \mathbf{S} + \tau_y^2 \mathbf{I}_p \quad \mathbf{K} + \sigma_z^2 \mathbf{I}_p = \mathbf{S} + \tau_z^2 \mathbf{I}_p.$$

For x-axis, we rewrite the equality as $\mathbf{K} - \mathbf{S} = (\sigma_x^2 - \tau_x^2)\mathbf{I}_p$. Since $\text{rank}(\mathbf{K}) < p/2$ and $\text{rank}(\mathbf{S}) < p/2$ as shown above, $\text{rank}(\mathbf{K} - \mathbf{S}) < p/2 + p/2 = p$. If $\sigma_x^2 \neq \tau_x^2$, then $\text{rank}((\sigma_x^2 - \tau_x^2)\mathbf{I}_p) = p$, which implies $\text{rank}(\mathbf{K} - \mathbf{S}) = p$ by the equality $\mathbf{K} - \mathbf{S} = (\sigma_x^2 - \tau_x^2)\mathbf{I}_p$, a contradiction. Therefore, $\sigma_x^2 = \tau_x^2$. By the same argument, we can show $\sigma_y^2 = \tau_y^2$ and $\sigma_z^2 = \tau_z^2$. Thus, the additional assumption that the axis variance matrices are low rank is sufficient to ensure identifiability.

#### 2.2 F-test for chromatin loop calling

Let $L_{jk}$ be the index set of all loop pairwise differences (values of the form $x_{ij} - x_{ik}$ for the x-axis and similarly for the other two axes; abbreviated as $x_{i,j,k}$ for clarity). For instance, if we are testing whether the 4th locus and the 8th locus form a loop, and there are four chromatin traces in total but the 3rd trace misses the 4th locus, then

$$L_{48} = \{(1, 4, 8), (2, 4, 8), (4, 4, 8)\}.$$

Similarly, let $B_{jk}$ denote the index set of background pairwise differences. We now describe the F-test for each axis. For the x-axis, we assume that after the filtering and normalization step, pairwise differences in $L_{jk}$ are independent and identically distributed as $\text{N}(0, \sigma^2_{\text{loop}})$ and pairwise differences in $B_{jk}$ are independent and identically distributed as $\text{N}(0, \sigma^2_{\text{bkgd}})$. The null that the $jk$th locus pair does not form a loop corresponds to $H_0 : \sigma^2_{\text{loop}} = \sigma^2_{\text{bkgd}}$ by the distance–variance relation explained in Methods.

To test this $H_0$, we derive the likelihood ratio test (LRT) statistic below. For simplicity, let $\sigma_1^2 = \sigma^2_{\text{loop}}$, $\sigma_2^2 = \sigma^2_{\text{bkgd}}$, $n_1 = |L_{jk}|$, $n_2 = |B_{jk}|$, $\{a_1, ..., a_{n_1}\} = \{x_{i,j,k} : (i, j, k) \in L_{jk}\}$, and $\{b_1, ..., b_{n_2}\} = \{x_{i,j,k} : (i, j, k) \in B_{jk}\}$. The model assumes $a_1, ..., a_{n_1} \sim_{\text{i.i.d.}} \text{N}(0, \sigma_1^2)$, $b_1, ..., b_{n_2} \sim_{\text{i.i.d.}} \text{N}(0, \sigma_2^2)$, and $(a_1, ..., a_{n_1})$ and $(b_1, ..., b_{n_2})$ are independent. The LRT statistic for $H_0 : \sigma_1^2 = \sigma_2^2$ is

$$L = \frac{\sup_{\sigma^2} f(a_1, ..., a_{n_1}|\sigma^2) f(b_1, ..., b_{n_2}|\sigma^2)}{\sup_{\sigma_1^2, \sigma_2^2} f(a_1, ..., a_{n_1}|\sigma_1^2) f(b_1, ..., b_{n_2}|\sigma_2^2)},$$

where $f(\cdot, ..., \cdot \mid \cdot)$ is the multivariate normal probability density function (PDF). The numerator is

$$\sup_{\sigma^2 \geq 0} (2\pi\sigma^2)^{-\frac{n_1+n_2}{2}} \exp\left( -\frac{1}{2\sigma^2} \left\{ \sum_{i=1}^{n_1} a_i^2 + \sum_{i=1}^{n_2} b_i^2 \right\} \right).$$

Taking the log and setting the derivative to zero gives

$$\frac{1}{2}(n_1 + n_2)\frac{1}{\sigma^2} = \frac{1}{2(\sigma^2)^2} \left\{ \sum_{i=1}^{n_1} a_i^2 + \sum_{i=1}^{n_2} b_i^2 \right\} \quad \Rightarrow \quad \hat{\sigma}^2 = \frac{1}{n_1 + n_2} \left\{ \sum_{i=1}^{n_1} a_i^2 + \sum_{i=1}^{n_2} b_i^2 \right\}.$$

The denominator is

$$\sup_{\sigma_1^2, \sigma_2^2 \geq 0} (2\pi\sigma_1^2)^{-\frac{n_1}{2}} \exp\left( -\frac{1}{2\sigma_1^2} \sum_{i=1}^{n_1} a_i^2 \right) (2\pi\sigma_2^2)^{-\frac{n_2}{2}} \exp\left( -\frac{1}{2\sigma_2^2} \sum_{i=1}^{n_2} b_i^2 \right).$$

By differentiating the log-likelihood and setting it to zero, we have

$$\hat{\sigma}_1^2 = \frac{1}{n_1}\sum_{i=1}^{n_1} a_i^2 \quad \text{and} \quad \hat{\sigma}_2^2 = \frac{1}{n_2}\sum_{i=1}^{n_2} b_i^2.$$

Therefore, the LRT statistic is

$$\begin{aligned}
L &= \left(\frac{\hat{\sigma}^2}{\hat{\sigma}_1^2}\right)^{-\frac{n_1}{2}} \left(\frac{\hat{\sigma}^2}{\hat{\sigma}_2^2}\right)^{-\frac{n_2}{2}} \exp\left(-\frac{1}{2\hat{\sigma}^2}\left\{\sum_{i=1}^{n_1} a_i^2 + \sum_{i=1}^{n_2} b_i^2\right\} + \frac{1}{2\hat{\sigma}_1^2}\sum_{i=1}^{n_1} a_i^2 + \frac{1}{2\hat{\sigma}_2^2}\sum_{i=1}^{n_2} b_i^2\right) \\
&= \left(\frac{\hat{\sigma}^2}{\hat{\sigma}_1^2}\right)^{-\frac{n_1}{2}} \left(\frac{\hat{\sigma}^2}{\hat{\sigma}_2^2}\right)^{-\frac{n_2}{2}} \exp\left(-\frac{1}{2}(n_1+n_2) + \frac{1}{2}n_1 + \frac{1}{2}n_2\right) \\
&= \left(\frac{\hat{\sigma}^2}{\hat{\sigma}_1^2}\right)^{-\frac{n_1}{2}} \left(\frac{\hat{\sigma}^2}{\hat{\sigma}_2^2}\right)^{-\frac{n_2}{2}}.
\end{aligned}$$

The LRT rejects $H_0$ if $L < k$, where $k$ controls the type I error $\alpha$ as $\alpha = \mathbb{P}(L < k \mid H_0)$. We further simplify $L$ to derive a closed-form expression. Since $(n_1+n_2)\hat{\sigma}^2 = n_1\hat{\sigma}_1^2 + n_2\hat{\sigma}_2^2$,

$$\begin{aligned}
L &= (n_1+n_2)^{\frac{1}{2}(n_1+n_2)} \left((n_1+n_2)\frac{\hat{\sigma}^2}{\hat{\sigma}_1^2}\right)^{-\frac{n_1}{2}} \left((n_1+n_2)\frac{\hat{\sigma}^2}{\hat{\sigma}_2^2}\right)^{-\frac{n_2}{2}} \\
&\propto \left(n_1 + \frac{n_2}{r}\right)^{-\frac{n_1}{2}} (n_2 + n_1 r)^{-\frac{n_2}{2}} \\
&\propto \left(n_1 + \frac{n_2}{r}\right)^{-n_1} (n_2 + n_1 r)^{-n_2},
\end{aligned}$$

where the first $\propto$ follows by dropping the constant $(n_1+n_2)^{\frac{1}{2}(n_1+n_2)}$ and the second $\propto$ follows by taking squares and noting the value is nonnegative. $r$ is defined as $r = \hat{\sigma}_1^2/\hat{\sigma}_2^2$. Therefore, $L < k$ for some $k$ is equivalent to $\left(n_1 + \frac{n_2}{r}\right)^{-n_1} (n_2 + n_1 r)^{-n_2} < k'$ for some $k'$.

We can make additional simplification by analyzing the function $f(r) = \left(n_1 + \frac{n_2}{r}\right)^{-n_1} (n_2 + n_1 r)^{-n_2}$. The first derivative of $f$ is

$$f'(r) = n_1\left(n_1 + \frac{n_2}{r}\right)^{-n_1-1} \frac{n_2}{r^2}(n_2+n_1 r)^{-n_2} - n_1 n_2\left(n_1 + \frac{n_2}{r}\right)^{-n_1} (n_2+n_1 r)^{-n_2-1},$$

which equals to zero if

$$\begin{aligned}
n_1 n_2 r^{-2}\left(n_1 + \frac{n_2}{r}\right)^{-n_1-1} (n_2+n_1 r)^{-n_2} &= n_1 n_2\left(n_1 + \frac{n_2}{r}\right)^{-n_1} (n_2+n_1 r)^{-n_2-1} \\
\iff \quad n_2 + n_1 r &= r^2\left(n_1 + \frac{n_2}{r}\right) \\
\iff \quad n_1 r^2 + (n_2 - n_1) r - n_2 &= 0,
\end{aligned}$$

i.e., $r = 1$ or $r = -n_2/n_1$. Since $r = \hat{\sigma}_1^2/\hat{\sigma}_2^2 > 0$, $f'(r) = 0$ if and only if $r = 1$. To see whether $f'(r) > 0$ for $r > 1$ or $r < 1$, take $r = n_2/n_1$. Then $f'(r) = (2n_1)^{-n_1-1}(2n_2)^{-n_2}\frac{n_1^2}{n_2^2} - (2n_1)^{-n_1}(2n_2)^{-n_2-1}$, and the ratio between the two terms is

$$(2n_1)^{-n_1-1}(2n_2)^{-n_2}\frac{n_1^2}{n_2^2}(2n_1)^{n_1}(2n_2)^{n_2+1} = \frac{n_2}{n_1}\frac{n_1^2}{n_2^2} = \frac{n_1}{n_2}$$

Thus, if $n_1 > n_2$, then $r < 1$ and $f'(r) > 0$; and if $n_2 > n_1$, then $r > 1$ and $f'(r) < 0$.

From the first derivative, $f'(r) > 0$ if $r < 1$ and $f'(r) < 0$ if $r > 1$. As a result, $f(r)$ increases as $r$ increases when $r \leq 1$, and $f(r)$ increases as $r$ decreases when $r > 1$. Therefore, the rejection rule $f(r) < k'$ is equivalent to

$$\begin{cases} r < k_1 & \text{if } r \leq 1 \\ r > k_2 & \text{if } r > 1 \end{cases}$$

for some $k_1, k_2$. Note $r$ has an $F$-distribution under $H_0 : \sigma_1^2 = \sigma_2^2 \equiv \sigma^2$ since

$$r = \frac{\hat{\sigma}_1^2}{\hat{\sigma}_2^2} = \frac{\frac{1}{n_1}\sum_{i=1}^{n_1} a_i^2}{\frac{1}{n_2}\sum_{i=1}^{n_2} b_i^2} = \frac{\frac{1}{n_1}\sum_{i=1}^{n_1} \frac{a_i^2}{\sigma^2}}{\frac{1}{n_2}\sum_{i=1}^{n_2} \frac{b_i^2}{\sigma^2}} =_d \frac{\frac{1}{n_1}\chi^2_{n_1}}{n_2 \chi^2_{n_2}} \sim F(n_1, n_2).$$

For loop calling, since the alternative is $H_1 : \sigma^2_{\text{loop}} < \sigma^2_{\text{bkgd}}$, we reject $H_0$ only if $\sigma^2_{\text{loop}}/\sigma^2_{\text{bkgd}} = r < k_1$. We can solve for $k_1$ for any pre-specified type I error rate $\alpha$ using the $F$-distribution:

$$\alpha = \mathbb{P}(\text{reject } H_0 | H_0) = \mathbb{P}(F(n_1, n_2) < k_1) \quad \Rightarrow \quad k_1 = F(n_1, n_2, \alpha),$$

where $F(n_1, n_2, \alpha)$ denote the $\alpha$-quantile of $F(n_1, n_2)$. The p-value is $p_x = \mathbb{P}(F(n_1, n_2) \leq r)$.

In summary, to test, for the x-axis, $H_0 : \sigma^2_{\text{loop}} = \sigma^2_{\text{bkgd}}$ versus $H_1 : \sigma^2_{\text{loop}} < \sigma^2_{\text{bkgd}}$, we compute

$$r = \frac{\hat{\sigma}^2_{\text{loop}}}{\hat{\sigma}^2_{\text{bkgd}}}, \quad \text{where} \quad \hat{\sigma}^2_{\text{loop}} = \frac{1}{|L_{jk}|} \sum_{(i,j,k) \in L_{jk}} x^2_{i,j,k} \quad \text{and} \quad \hat{\sigma}^2_{\text{bkgd}} = \frac{1}{|B_{jk}|} \sum_{(i,j,k) \in B_{jk}} x^2_{i,j,k}$$

and calculate the p-value $p_x$. This procedure is then repeated for the y and z axes, and the final p-value for the null that the $jk$ locus pair is not a loop is obtained by $p = w_x \tan\{(0.5 - p_x)\pi\} + w_y \tan\{(0.5 - p_y)\pi\} + w_z \tan\{(0.5 - p_z)\pi\}$.

##### 2.3 F-test for TAD calling

Suppose we are testing whether the $i$th locus is a TAD boundary. Let $E_i$ and $A_i$ be the index sets for inter-region pairwise differences and intra-region pairwise differences. For example, we are testing the 6th locus with a window size of two loci, and there are two chromatin traces with no missing data. Then the index sets are

$$\begin{aligned} E_i = \{&(1,4,7), (1,5,7), (1,4,8), (1,5,8) \\ &(2,4,7), (2,5,7), (2,4,8), (2,5,8)\} \\ A_i = \{&(1,4,5), (1,4,6), (1,5,6), (1,6,7), (1,6,8), (1,7,8) \\ &(2,4,5), (2,4,6), (2,5,6), (2,6,7), (2,6,8), (2,7,8)\}. \end{aligned}$$

This is illustrated below:

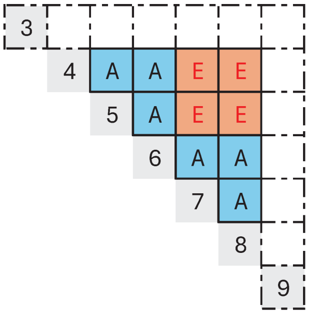

As in chromatin loop calling, pairwise differences in $E_i$ and $A_i$ are independent and identically distributed as $\mathrm{N}(0, \sigma^2_{\text{inter}})$ and $\mathrm{N}(0, \sigma^2_{\text{intra}})$. For each axis, the corresponding null and alternative hypotheses are $H_0 : \sigma^2_{\text{inter}} = \sigma^2_{\text{intra}}$ and $H_1 : \sigma^2_{\text{inter}} > \sigma^2_{\text{intra}}$.

Using the same argument, for x-axis, the test statistic is

$$r = \frac{\hat{\sigma}^2_{\text{intra}}}{\hat{\sigma}^2_{\text{inter}}}, \quad \text{where} \quad \hat{\sigma}^2_{\text{intra}} = \frac{1}{|A_i|} \sum_{(i',j,k)\in A_i} x^2_{i',j,k} \quad \text{and} \quad \hat{\sigma}^2_{\text{inter}} = \frac{1}{|E_i|} \sum_{(i',j,k)\in E_i} x^2_{i',j,k},$$

and the p-value is $p_x = \mathbb{P}(F(|A_i|, |E_i|) \leq r)$. After testing each axis, the final p-value is obtained by Cauchy combination test as before.

#### 2.4 Test statistics computation based on axis variance matrix

The test statistics derived in the previous two subsections involve summing over subsets of pairwise difference matrices (i.e., $x_{i,j,k}$, $y_{i,j,k}$, and $z_{i,j,k}$), which take $O(np^2)$ space to store. Thus, instead of storing the entire pairwise difference matrices, we store only the axis variance matrix (a $p \times p$ matrix for each axis) and a corresponding count matrix (the same shape as the variance matrix).

Take the x-axis as an example. Let $I_{jk}$ denote the index set for the $jk$th pairwise differences. If there are four traces with the second trace missing the 25 pair, then

$$I_{25} = \{(1, 2, 5), (3, 2, 5), (4, 2, 5)\}.$$

The $jk$th entry of the count matrix is defined as $C_{jk} = |I_{jk}|$, i.e., the number of available pairwise differences. For x axis, the $jk$th entry of the axis variance matrix is defined as

$$V^{(x)}_{jk} = \frac{1}{|I_{jk}|} \sum_{(i,j,k)\in I_{jk}} (x_{ij} - x_{ik})^2.$$

For loop calling, the test statistic for the $jk$th locus pair is $r = \hat{\sigma}^2_{\text{loop}}/\hat{\sigma}^2_{\text{bkgd}}$, where $\hat{\sigma}^2_{\text{loop}} = V^{(x)}_{jk}$ and

$$\hat{\sigma}^2_{\text{bkgd}} = \frac{1}{|B_{jk}|} \sum_{(i,j,k)\in B_{jk}} (x_{ij} - x_{ik})^2 = \frac{1}{\sum_{j'k'} C_{j'k'}} \sum_{j'k'} C_{j'k'} V^{(x)}_{j'k'},$$

where the summation is taken over all $j'k'$ pairs within the local background. Therefore, for loop calling, the test statistic can be computed only using the axis variance matrix and the count matrix. For TAD calling, the test statistic of the $j$th locus is $r = \hat{\sigma}^2_{\text{intra}}/\hat{\sigma}^2_{\text{inter}}$. Note

$$\hat{\sigma}^2_{\text{intra}} = \frac{1}{|A_j|} \sum_{(i,j',k')\in A} (x_{ij'} - x_{ik'})^2 = \frac{1}{\sum_{j'k'} C_{j'k'}} \sum_{j'k'} C_{j'k'} V^{(x)}_{j'k'}$$

where the summation is taken over all intra-region $j'k'$ pairs. Similarly, $\hat{\sigma}^2_{\text{inter}}$ can be calculated using only the axis variance matrix and the count matrix.

#### 2.5 Second eigenvector for A/B compartment calling

As mentioned in Methods, we model A/B compartment by an SBM: each imaged locus has an underlying membership, either A or B, and loci with the same membership are spatially closer to each other. In this subsection, we first explain the reason for using the second eigenvector and then describe the spectral clustering algorithm.

We assume loci from the same compartment have a probability of $p$ to interact with each other, and loci from different compartments have a probability of $q$ with $q < p$ to interact with each other. We define $\mathbf{W} \in \mathbb{R}^{2\times 2}$ and the membership matrix $\mathbf{C} \in \mathbb{R}^{p\times 2}$ as

$$\mathbf{W} = \begin{bmatrix} p & q \\ q & p \end{bmatrix}, \quad C_{i1} = \begin{cases} 1 & i\text{th locus is A} \\ 0 & i\text{th locus is B} \end{cases} \quad \text{and} \quad C_{i2} = \begin{cases} 1 & i\text{th locus is B} \\ 0 & i\text{th locus is A} \end{cases}.$$

The adjacency matrix $\mathbf{P} \in \mathbb{R}^{p\times p}$ is defined so that $P_{ij}$ equals the probability that the $i$th locus and the $j$th locus interact with each other. Note $\mathbf{P} = \mathbf{C}\mathbf{W}\mathbf{C}^\top$ since the $ij$th entry of $\mathbf{C}\mathbf{W}\mathbf{C}^\top$ is

$$\begin{pmatrix} C_{i1} & C_{i2} \end{pmatrix} \begin{bmatrix} p & q \\ q & p \end{bmatrix} \begin{pmatrix} C_{j1} \\ C_{j2} \end{pmatrix} = \begin{cases} p & C_{i1} = C_{j1} \\ q & C_{i1} \neq C_{j1} \end{cases} = \begin{cases} p & \text{same compartment} \\ q & \text{different compartment} \end{cases} = P_{ij}.$$

We now explain how locus membership relates to the second eigenvector of $\mathbf{P}$. Essentially, we will show that the $i$th value and the $j$th value of the second eigenvector are the same if the $i$th and the $j$th loci belong to the same compartment, and the values are different if their memberships differ. Therefore, the second eigenvector completely determines locus membership.

Let $p_1$ and $p_2$ denote the number of A loci and the number of B loci, respectively. Define the diagonal matrix $\mathbf{D} = \operatorname{diag}(p_1, p_2)$. Then

$$\mathbf{P} = \mathbf{C}\mathbf{W}\mathbf{C}^\top = \mathbf{C}\mathbf{D}^{-\frac{1}{2}}\mathbf{D}^{\frac{1}{2}}\mathbf{W}\mathbf{D}^{\frac{1}{2}}\mathbf{D}^{-\frac{1}{2}}\mathbf{C}^\top = \mathbf{C}\mathbf{D}^{-\frac{1}{2}}\mathbf{V}\boldsymbol{\Lambda}\mathbf{V}^\top\mathbf{D}^{-\frac{1}{2}}\mathbf{C}^\top := \mathbf{U}\boldsymbol{\Lambda}\mathbf{U}^\top,$$

where $\mathbf{V}\boldsymbol{\Lambda}\mathbf{V}^\top$ is the eigenvalue decomposition of $\mathbf{D}^{\frac{1}{2}}\mathbf{W}\mathbf{D}^{\frac{1}{2}}$. Note $\mathbf{U}\boldsymbol{\Lambda}\mathbf{U}^\top$ is the eigenvalue decomposition of $\mathbf{P}$ since $\boldsymbol{\Lambda}$ is diagonal and $\mathbf{U}$ is orthonormal as

$$\mathbf{U}^\top\mathbf{U} = \mathbf{V}^\top\mathbf{D}^{-\frac{1}{2}}\mathbf{C}^\top\mathbf{C}\mathbf{D}^{-\frac{1}{2}}\mathbf{V} = \mathbf{V}^\top \begin{bmatrix} p_1^{-\frac{1}{2}} & 0 \\ 0 & p_2^{-\frac{1}{2}} \end{bmatrix} \begin{bmatrix} p_1 & 0 \\ 0 & p_2 \end{bmatrix} \begin{bmatrix} p_1^{-\frac{1}{2}} & 0 \\ 0 & p_2^{-\frac{1}{2}} \end{bmatrix} \mathbf{V} = \mathbf{V}^\top\mathbf{I}\mathbf{V} = \mathbf{I}.$$

We now derive the second eigenvector (the second column of $\mathbf{U}$). This is achieved by deriving $\mathbf{V}$ and using $\mathbf{U} = \mathbf{C}\mathbf{D}^{-\frac{1}{2}}\mathbf{V}$. The eigenvalues are determined by solving $\det(\mathbf{D}^{\frac{1}{2}}\mathbf{W}\mathbf{D}^{\frac{1}{2}} - \lambda\mathbf{I}) = 0$:

$$\mathbf{D}^{\frac{1}{2}}\mathbf{W}\mathbf{D}^{\frac{1}{2}} - \lambda\mathbf{I} = \begin{bmatrix} p_1 p - \lambda & \sqrt{p_1 p_2} q \\ \sqrt{p_1 p_2} q & p_2 p - \lambda \end{bmatrix} \quad \Rightarrow \quad (p_1 p - \lambda)(p_2 p - \lambda) = p_1 p_2 q^2,$$

which is $\lambda^2 - \lambda(p_1 + p_2)p + p_1 p_2(p^2 - q^2) = 0$ and has solutions

$$\lambda = \frac{1}{2}\left\{(p_1 + p_2)p \pm \sqrt{(p_1 - p_2)^2 p^2 + 4 p_1 p_2 q^2}\right\}.$$

Let $\lambda_1$ and $\lambda_2$ denote the two eigenvalues with $\lambda_1 > \lambda_2$. The second eigenvector of $\mathbf{D}^{\frac{1}{2}}\mathbf{W}\mathbf{D}^{\frac{1}{2}}$ (i.e., the second column of $\mathbf{V}$) satisfies

$$\mathbf{D}^{\frac{1}{2}}\mathbf{W}\mathbf{D}^{\frac{1}{2}}\mathbf{v} = \lambda_2\mathbf{v} \quad \Rightarrow \quad \begin{cases} p_1 p v_1 + \sqrt{p_1 p_2} q v_2 = \lambda_2 v_1 \\ \sqrt{p_1 p_2} q v_1 + p_2 p v_2 = \lambda_2 v_2 \end{cases} \quad \Rightarrow \quad v_1 = \frac{\sqrt{p_1 p_2} q}{\lambda_2 - p_1 p} v_2.$$

Thus, the $i$th entry of the second eigenvector of $\mathbf{P}$ is

$$\begin{pmatrix} C_{i1} & C_{i2} \end{pmatrix} \begin{bmatrix} p_1^{-\frac{1}{2}} & 0 \\ 0 & p_2^{-\frac{1}{2}} \end{bmatrix} \begin{pmatrix} v_1 \\ v_2 \end{pmatrix} = \begin{pmatrix} C_{i1} & C_{i2} \end{pmatrix} \begin{pmatrix} p_1^{-\frac{1}{2}} v_1 \\ p_2^{-\frac{1}{2}} v_2 \end{pmatrix} = \begin{cases} p_1^{-\frac{1}{2}} v_1 & i\text{th locus is A} \\ p_2^{-\frac{1}{2}} v_2 & i\text{th locus is B} \end{cases}.$$

Thus, to show the claim that the $i$th value and $j$th value of the second eigenvector are the same if and only if the $i$th and the $j$th loci belong to the same compartment, it suffices to show $p_1^{-\frac{1}{2}}v_1 \neq p_2^{-\frac{1}{2}}v_2$. Note $p_1^{-\frac{1}{2}}v_1 = p_2^{-\frac{1}{2}}v_2$ if and only if

$$\begin{aligned} p_1^{-\frac{1}{2}} \frac{\sqrt{p_1 p_2} q}{\lambda_2 - p_1 p} v_2 = p_2^{-\frac{1}{2}} v_2 \quad &\iff \quad p_2 q = \lambda_2 - p_1 p \\ &\iff \quad p_2 q + p_1 p - \frac{1}{2}(p_1 + p_2)p + \frac{1}{2}\sqrt{(p_1 - p_2)^2 p^2 + 4p_1 p_2 q^2} = 0 \\ &\iff \quad p_2 q + \frac{1}{2}\left\{(p_1 - p_2)p + \sqrt{(p_1 - p_2)^2 p^2 + 4p_1 p_2 q^2}\right\} = 0. \end{aligned}$$

The term in the brackets is

$$\begin{aligned} &(p_1 - p_2)p + \sqrt{(p_1 - p_2)^2 p^2 + 4p_1 p_2 q^2} \\ &> (p_1 - p_2)p + \sqrt{(p_1 - p_2)^2 p^2} \qquad \text{since } 4p_1 p_2 q^2 > 0 \text{ and } \sqrt{\cdots} \text{ is increasing} \\ &= (p_1 - p_2)p + |p_1 - p_2|p \geq 0. \end{aligned}$$

As both $p_2 q$ and the term in the brackets are greater than 0, $p_1^{-\frac{1}{2}}v_1 \neq p_2^{-\frac{1}{2}}v_2$.

In contrast, the first eigenvector of $\mathbf{P}$ does not guarantee the above property. Specifically, when $p_1 = p_2 \equiv p_0$, the first eigenvalue is

$$\lambda_1 = \frac{1}{2}\left\{2p_0 p + \sqrt{(p_0 - p_0)^2 p^2 + 4p_0 p_0 q^2}\right\} = p_0 p + p_0 q = p_0(p + q),$$

and the corresponding eigenvector of $\mathbf{D}^{\frac{1}{2}}\mathbf{W}\mathbf{D}^{\frac{1}{2}}$ is $v_1 = \frac{\sqrt{p_0 p_0} q}{\lambda_1 - p_0 p} v_2 = \frac{p_0 q}{p_0(p+q) - p_0 p} v_2 = v_2$. Thus

$$\begin{pmatrix} C_{i1} & C_{i2} \end{pmatrix} \begin{bmatrix} p_0^{-\frac{1}{2}} & 0 \\ 0 & p_0^{-\frac{1}{2}} \end{bmatrix} \begin{pmatrix} v_1 \\ v_2 \end{pmatrix} = \begin{pmatrix} C_{i1} & C_{i2} \end{pmatrix} \begin{pmatrix} p_0^{-\frac{1}{2}} v_1 \\ p_0^{-\frac{1}{2}} v_2 \end{pmatrix} = p_0^{-\frac{1}{2}} v_1.$$

As the first eigenvector of $\mathbf{P}$ is a constant, it cannot distinguish the two compartments.

We now describe the spectral clustering algorithm. Although the adjacency matrix $\mathbf{P}$ is not observed, it can be approximated by the similarity matrix calculated from the observed data. As the second eigenvector of $\mathbf{P}$ corresponds to locus membership, we expect that the second eigenvector of the similarity matrix have a similar property. However, since the eigenvalue decomposition is not performed directly on $\mathbf{P}$, the second eigenvector typically consists of more than two values, so K-means with $K = 2$ is applied to the second eigenvector, and the two clusters found are treated as the compartment assignment.

As measurement error differs for the x, y, and z axes, we create a similarity matrix for each axis. For example, the $jk$th entry of the similarity matrix for the x-axis is

$$\exp\left\{-\frac{1}{n}\sum_{i=1}^{n}(x_{ij} - x_{ik})^2\right\},$$

where $x_{ij} - x_{ik}$ is filtered and normalized as described in Methods. If there are missing values, the summation is taken over available pairwise differences, and the denominator is adjusted accordingly. Then eigenvalue decomposition is performed on each similarity matrix, and the second eigenvector is retained. This creates three $p$-dimensional vectors, forming the three-dimensional feature space described in Methods, and we employ K-means on this space to assign A/B compartments.
